# CONSTANS alters the circadian clock in *Arabidopsis thaliana*

**DOI:** 10.1101/2023.01.19.524697

**Authors:** Pedro de los Reyes, Francisco J Romero-Campero, He Gao, Gloria Serrano-Bueno, Jose M Romero, Federico Valverde

## Abstract

Plants are sessile organisms that have acquired highly plastic developmental strategies to adapt to the environment. Among these processes, the floral transition is essential to ensure reproductive success and is finely regulated by several internal and external genetic networks. The photoperiodic pathway, which controls the plant response to day length, is one of the most important pathways controlling flowering. In *Arabidopsis* photoperiodic flowering, *CONSTANS* (*CO*) is the central gene activating the expression of the florigen *FLOWERING LOCUS T* (*FT)* in the leaves at the end of a long day. *CO* expression is strongly regulated by the circadian clock. However, to date, no evidence has been reported regarding a feedback loop from the photoperiod pathway back to the circadian clock. Using transcriptional networks, we have identified relevant network motifs regulating the interplay between the circadian clock and the photoperiod pathway. Gene expression, chromatin immunoprecipitation experiments and phenotypic analysis allowed us to elucidate the role of CO over the circadian clock. Plants with altered *CO* expression showed a different internal clock period, measured by daily rhythmic movements in the leaves. We show that CO is able to activate key genes related to the circadian clock, such as *CCA1*, *LHY*, *PRR5* and *GI,* at the end of a long day by binding to specific sites on their promoters. Moreover, a significant number of PRR5 repressed target genes are upregulated by CO, and this could explain the phase transition promoted by CO. The CO-PRR5 complex interacts with the bZIP transcription factor HY5 and helps to localize the complex in the promoters of clock genes. Our results indicate that there may be a feedback loop in which CO communicates back to the circadian clock, feeding seasonal information to the circadian system.

## Introduction

In order to interact with a regularly changing environment, plants have developed a great variety of morphological and physiological responses. The transcriptional programs that control these responses allow plants to respond to external stimuli with a high degree of plasticity throughout their life cycle (De Mendoza et al., 2013). One of the most important decisions in the plant life cycle is flowering, which activates the molecular and metabolic changes that induce the transition from the vegetative to the reproductive growth programs (Piñeiro and Jarillo, 2013; Andrés and Coupland, 2010; Kinoshita and Richter, 2020). This transition is finely regulated by multiple factors, since making this decision at the right moment ensures reproductive and evolutionary success. Thus, plants are able to recognize seasonal changes by measuring day length (photoperiod), allowing them to adapt to and grow at different latitudes. The photoperiodic pathway is one of the major components controlling flowering and is regulated by light sensors and an autonomous internal timekeeper called the circadian clock (Sanchez and Kay, 2016; Shim et al., 2017). The central gene of the photoperiod pathway is *CONSTANS* (*CO*), which acts as integrator of day length signals and is an output of the clock (Putterill et al., 1995; Suárez-López et al., 2001; Valverde, 2011).

*CO* encodes a protein with two conserved protein domains: two B-boxes located at the amino terminus, involved in protein-protein interactions, and a carboxyl CCT (CONSTANS, CO LIKE, TIMING OF CAB1) domain, which includes a nuclear transport signal and mediates interactions with DNA and diverse proteins (Laubinger et al., 2006; Valverde, 2011). Recently, a role for the central part of CO in regulating floral senescence has been proposed (Serrano-Bueno et al., 2022). *CO* is a hub in the photoperiodic regulatory network; it receives signals from a multitude of different pathways, such as day length and circadian clock (Serrano-Bueno et al., 2021), and triggers the expression of the florigen *FLOWERING LOCUS T* (*FT*), which induces flower differentiation (Ahn et al, 2008). The circadian clock is an endogenous transcriptional clock that produces biological rhythms with periods of 24 h. This inner clock allows plants to synchronize and anticipate external changes and is one of the most important factors regulating flowering. In *Arabidopsis*, the circadian clock consists of three transcriptional loops, the core loop (CCA1, LHY and TOC1), the morning loop (PRR5, PRR7 and PRR9) and the evening loop (LUX, ELF3, ELF4 and GIGANTEA) (McClung, 2006; Sanchez and Kay, 2016). *CO* expression and protein stability are highly regulated by the circadian clock and light inputs (Valverde et al, 2004; Suárez-López et al., 2001; Mizoguchi et al., 2005). The GIGANTEA (GI) and FLAVIN-BINDING, KELCH REPEAT, F-BOX 1 (FKF1) complex participates in this control by regulating *CO* transcription by binding to its promoter in the late afternoon and promoting the degradation of CYCLING DOF FACTORS (CDFs), which act as repressors of *CO* expression (Imaizumi et al., 2005; Sawa et al., 2007). Later, FLOWERING BHLH proteins (FBHs) can bind to these genomic sites in the *CO* promoter to activate its expression (Ito et al., 2012). In addition, The PSEUDO RESPONSE REGULATORs PRR9, PRR7 and PRR5 indirectly promote photoperiodic flowering by repressing *CDF1* (Nakamichi et al., 2007).

Although the *CO* transcript is highly expressed from late afternoon to dawn, in terms of protein stability, CO is unstable in the dark and stable during the daytime via a process mediated by different photoreceptors and E3 ubiquitin ligases (Valverde et al., 2004). During the daytime, the RING finger-containing E3 ubiquitin ligase HOS1 (HIGH EXPRESSION OF OSMOTICALLY RESPONSIVE GENES1) interacts with CO and targets it for degradation (Lazaro et al., 2012), thus allowing activation of *FT* at the right moment of the day. During the night time, COP1 and SPA1 promote CO degradation (Laubinger et al., 2006; Jang et al., 2008). After dawn, several photoreceptors begin to stabilize CO to promote its accumulation during the first hours of the day. In response to blue light, CRY1 and CRY2 interact with COP1 and SPA1 to avoid the degradation of CO by the COP1-SPA1 complex (Yanovsky and Kay, 2002; Liu et al., 2008; Liu et al., 2011; Zuo et al., 2011). Moreover, during the day, two phytochromes affect CO stability in different ways; in response to red light, PHYB interacts with HOS1 to promote the degradation of CO, but in the late afternoon, PHYA is able to disrupt the COP1-SPA1 complex to stabilize CO in presence of far-red light (Valverde et al., 2004, Lazaro et al., 2012; Endo et al., 2013; Hajdu et al., 2015; Lazaro et al., 2015; Sheerin et al., 2015).

In addition to the described photoreceptor-pathways, there are some circadian-related proteins that regulate CO accumulation in response to blue light. In the morning, ZEITLUPE (ZTL) forms a complex with GI to inhibit CO stability (Song et al., 2014), but in the afternoon, blue-light enhances CO accumulation by interacting with FKF1 (Song et al., 2012). In fact, FKF1, GI and CO could form a complex that stabilizes CO (Song et al., 2014). Recently, it has been described that PRRs stabilize CO and promote flowering in response to day length (Hayama et al., 2017). CO stability and its role as transcriptional regulator are both affected. DELLA proteins, the immunophilin FKBP12, the RING domain protein BOI, the B-boxes BBX19 and miP1a, and the chromatin remodeling factor PICKLE (PKL) can regulate CO transcriptional function in different ways (Wang et al., 2014; Nguyen et al., 2015; Graeff et al., 2016; Wang et al., 2016; Xu et al., 2016; Jing et al., 2019; Serrano-Bueno et al., 2020)). Therefore, CO function is highly regulated by multiple pathways, all of them ensuring the precise activation of *FT* and the initiation of flowering. Regarding other biological processes regulated by CO, it has been described that CO is able to modify starch metabolism through the activation of the *GRANULE BOUND STARCH SYNTHASE* (*GBSS*) gene (Ortiz-Marchena et al., 2014). CO may also be involved in stomatal opening (Ando et al., 2013) and play a role in floral senescence (Serrano-Bueno et al., 2022). However, the involvement of CO in other physiological processes is still largely unknown.

Although considered an output of the circadian machinery, it is not known if CO is able to affect the clock. However, since CO presents a CCT domain in its carboxyl terminus, similar to that present in PRRs, that may allow binding to circadian genes promoters (Nakamichi et al., 2010), it was suggested that CO could participate in circadian clock control (Strayer et al., 2000). Circadian network retrograde signaling is often found, and in this study, we provide conclusive evidence that in addition to being an output of the clock, CO can also regulate the expression of key genes involved in the circadian time pacer. Therefore, the transcriptional programs controlling the floral transition and the circadian clock are intertwined, constituting a complex system exhibiting feedback loops. Recently, in order to unravel this system, omics techniques based on high throughput sequencing have been applied. However, these data are commonly fragmented, and molecular systems biology approaches are necessary to integrate them and obtain a global comprehension of the interactions between the transcriptional programs controlling the floral transition and the circadian clock.

Network theory constitutes one of the central paradigms in data integration and analysis in molecular systems biology (Barabási, 2015). The integration of massive amounts of omics data has helped generate biological networks for photosynthetic organisms that reveal emergent properties and improve the understanding of processes such as the evolution of CO-like transcription factors (TFs) in *Viridiplantae* (Romero-Campero et al., 2013), the conservation of circadian patterns across the plant phylogenetic tree (de los Reyes et al., 2017), carbon mobilization during the floral transition in *Arabidopsis* (Ortiz-Marchena et al., 2014) and the involvement of hormonal control (Serrano-Bueno et al., 2022). In this work, we have constructed a transcriptional gene network by integrating deep sequencing of chromatin immunoprecipitation (ChIP) data generated from key TFs involved in the floral development/transition and circadian clock control. Feeding data on CO into the analysis of this network allowed us to propose a new function for photoperiodic signaling: the formation of a feed-back loop regulating the circadian clock in *Arabidopsis*.

## Results

### Transcriptional networks reveal a connection between the circadian clock and photoperiodic signaling

To examine the interplay between the transcriptional programs controlling the circadian clock and photoperiod, we analyzed ChIP-seq data from 33 Arabidopsis TFs involved in light signaling, flowering and the circadian clock and built a transcriptional network called CircadianFloralNet. This network is composed of 20,601 nodes and 89,377 edges, comprising 75% of the *Arabidopsis* genome (Figure S1A). We also generated from CircadianFloralNet a transcriptional gene network containing only the analyzed TFs and their interactions (Figure S1B). This network constitutes the regulator core of CircadianFloralNet and was called CircadianFloralTFNet. Therefore, CircadianFloralTFNet is a transcriptional gene network that captures the transcriptional program governing the interaction between the circadian clock, light and the flowering process. CircadianFloralNet is a scale-free network showing a power-law node degree distribution (Figure S1C) according to the Kolmogorov-Smirnov test (Goldstein et al., 2004). Furthermore, it is a small-world network, implying that the distance between two random nodes is shorter than expected (Figure S1D). These topological properties indicate that transcriptional programs governing these processes are highly connected (Wagner and Fell., 2001; Tong et al., 2004; Yook et al., 2004).

To determine the possible input of photoperiod in this network, we carried out a genome-wide analysis of chromatin immunoprecipitation sequencing (ChIP-seq) experiments in 35S:*CO* and *co-10* 7-DAG seedlings around *Zeitgeber* Time (ZT) 14 under LD using specific CO antibodies (Serrano-Bueno et al., 2020). The analysis of CO ChIP-seq data identified 3,214 peaks and 2,418 putative target genes (Table S1), and we fed this information into CircadianFloralNet. Next, we generated CircadianFloralTFNet including CO and classified the nodes into light, flowering or clock genes based on their TAIR classifications (Figure 1A). CO was found to bind to the promoters of the *PIF1*, *PIF3* and *PIF4* genes related to light signals and to the promoters of *SOC1* and *SVP*, which are specifically associated with flowering time control. However, all genes clustering in the clock function in CircadianFloralTFNet, including *CCA1*, *LHY*, *TOC1, PRR5, PRR7,* and *PRR9,* showed one or several peaks of CO binding on their promoters (Figure 1A), indicating that CO significantly bound to the promoters of genes involved in circadian rhythm regulation, including the core components of the clock, *CCA1*/*LHY* and *TOC1*.

**Figure 1.**
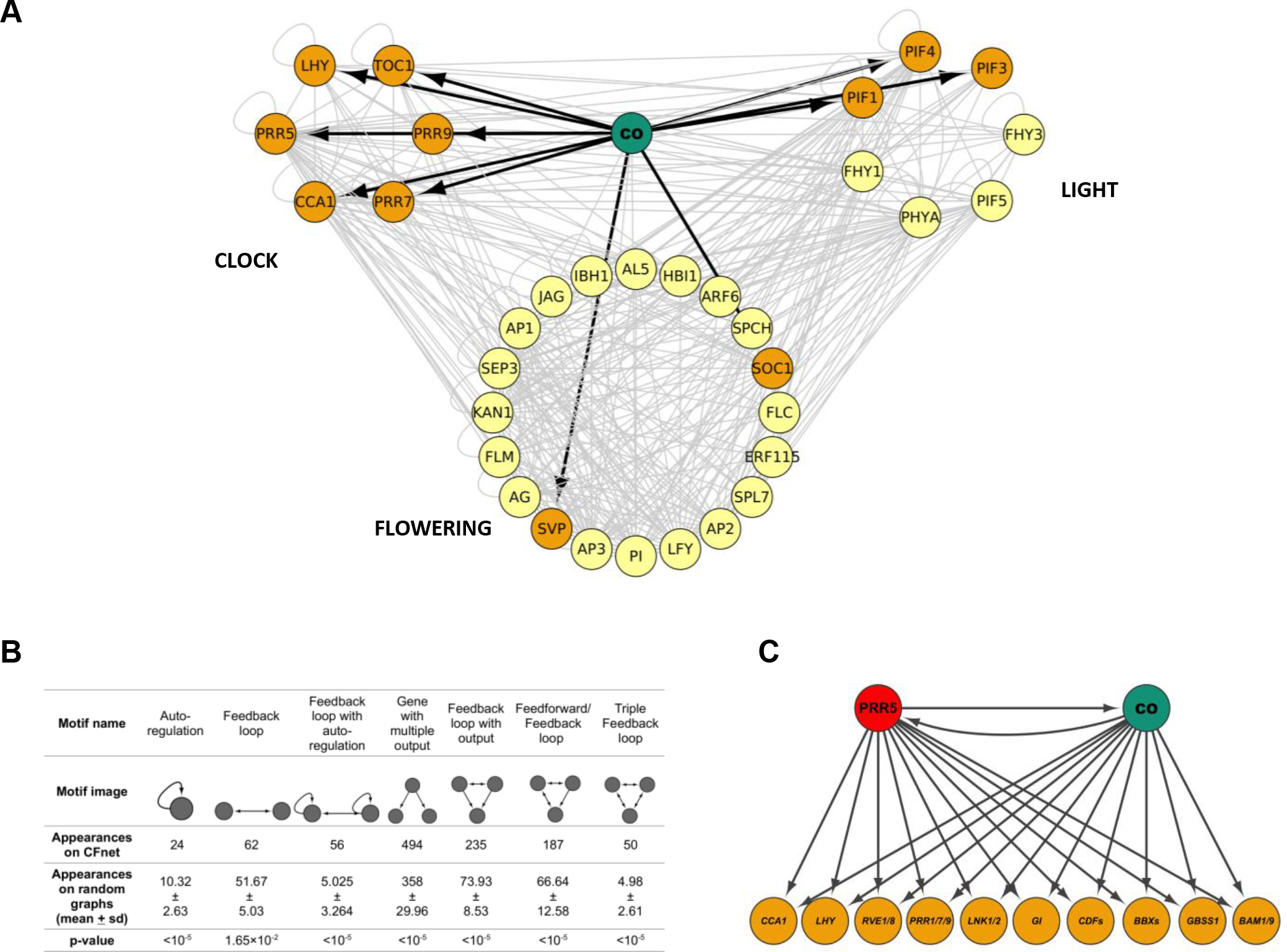
**CONSTANS regulates key circadian clock-related genes in the transcriptional network CircadianFloralNet**. (A) Graphical representation of CircadianFloralTFNet, a derived network including only the analyzed TFs and their interactions. Nodes are clustered in CLOCK, LIGHT or FLOWERING. CO target genes are colored in dark orange. (B) Most representative network motifs found in CircadianFloralTFNet showing the significance (in p-values) of their appearances. (C) Feedback loop with multiple outputs containing CO and PRR5 as regulators with a set of common target genes involved in the circadian clock and photoperiodic signaling.

In order to identify relevant non-random components in the interactions between the circadian clock and photoperiod, we performed a network motif analysis on CircadianFloralTFNet. A network motif is a subgraph that appears in the network of interest at a significantly higher number of times when compared to similar random networks. Therefore, we tested the significance of all possible subgraphs consisting of one, two or three nodes by generating 100,000 random graphs with the same topological features as the original network. The most represented motifs identified are shown in Figure 1B. One of the most significant and common motifs was the ‘feedback loop with output’ (p-value <10^-5^), which meant that there is a high interconnection among the transcriptional programs analyzed.

The connectivity we identified in CircadianFloralNet confirmed the results of previous studies (Suárez-López et al., 2001; Imaizumi et al., 2005; Sawa et al., 2007) that established that *CO* expression was regulated by the circadian clock. Specifically, PRR5 binds to cis elements in the *CO* promoter in order to regulate its expression, and consequently, both genes are connected in CircadianFloralNet. Additionally, a substantial number of genes of the *CO*-like (*COL*) family are regulated by PRR5 and other central proteins of the circadian clock, such as *CCA1* and *PRR7*, as shown in the subnetwork presented in Figure S1E. Since the feedback loop is a network motif in CircadianFloralTFNet and PRR5 regulates *CO,* it was expected that CO, in turn, would also regulate the expression of *PRR5*. We verified CO binding to the *PRR5* promoter through ChIP-seq and ChIP-QPCR experiments (Figure S1F and S4B). This result implies that one of the interactions between the flowering program and the circadian clock is exerted by CO and PRR5.

In fact, based on ChIP-seq data, CO and PRR5 constitute a ‘feedback loop with multiple output’ motif, represented in Figure 1C, that regulates a set of common target genes related with the circadian clock (*CCA1, LHY, RVE1,8, TOC1, PRR7,9, LNK1,2*) and photoperiod signaling (*GI, CDFs, BBXs, GBBS1 and BAM1,9*).

### *CONSTANS* alters the expression of circadian clock genes

To investigate the proposed effect of CO on the clock, we carried out RNA-seq experiments from different plants: general expression (35S:*CO*), phloem-specific, where CO floral function takes place, (SUC2:*CO*) and Col-0 (WT) (GSE236178). We extracted RNA from LD (LD, 16 h light: 8 h night) grown seedlings collected in the evening at ZT14, where CO activity is at a peak (Valverde et al., 2004). Similarly, we included an available RNA-seq dataset of Col-0 and *co-10* mutant plants grown in continuous light (LL) (Gnesutta et al., 2017; GSE205675). After analyzing the data via standard pipelines (see methods), we found 637 and 1765 upregulated genes in 35S:*CO* and SUC2:*CO* compared to WT, respectively, and 714 downregulated genes in *co-10* plants (Table S2). To obtain high-fidelity *CO*-regulated genes, we overlapped the three sets of genes in a Venn diagram consisting of 70 intersecting genes (Figure 2A, Table S3). According to the GO-term enrichment analysis, these genes are involved in several biological processes, such as the response to light stimuli or photoperiodism and, and as we expected, rhythmic processes and circadian rhythm with the highest probability (Figure 2A, Table S4). Specifically, several key circadian clock genes (*CCA1*, *LHY*, *GI*, *PRR9* and *LNK1*) and clock-output genes (*CDF1*, *CDF6*, *BBX28*, *BBX29*, *BBX30* and *BBX32*) showed higher expression in SUC2:*CO* plants and lower expression in *co-10* mutants (Figures 2C, S2).

**Figure 2.**
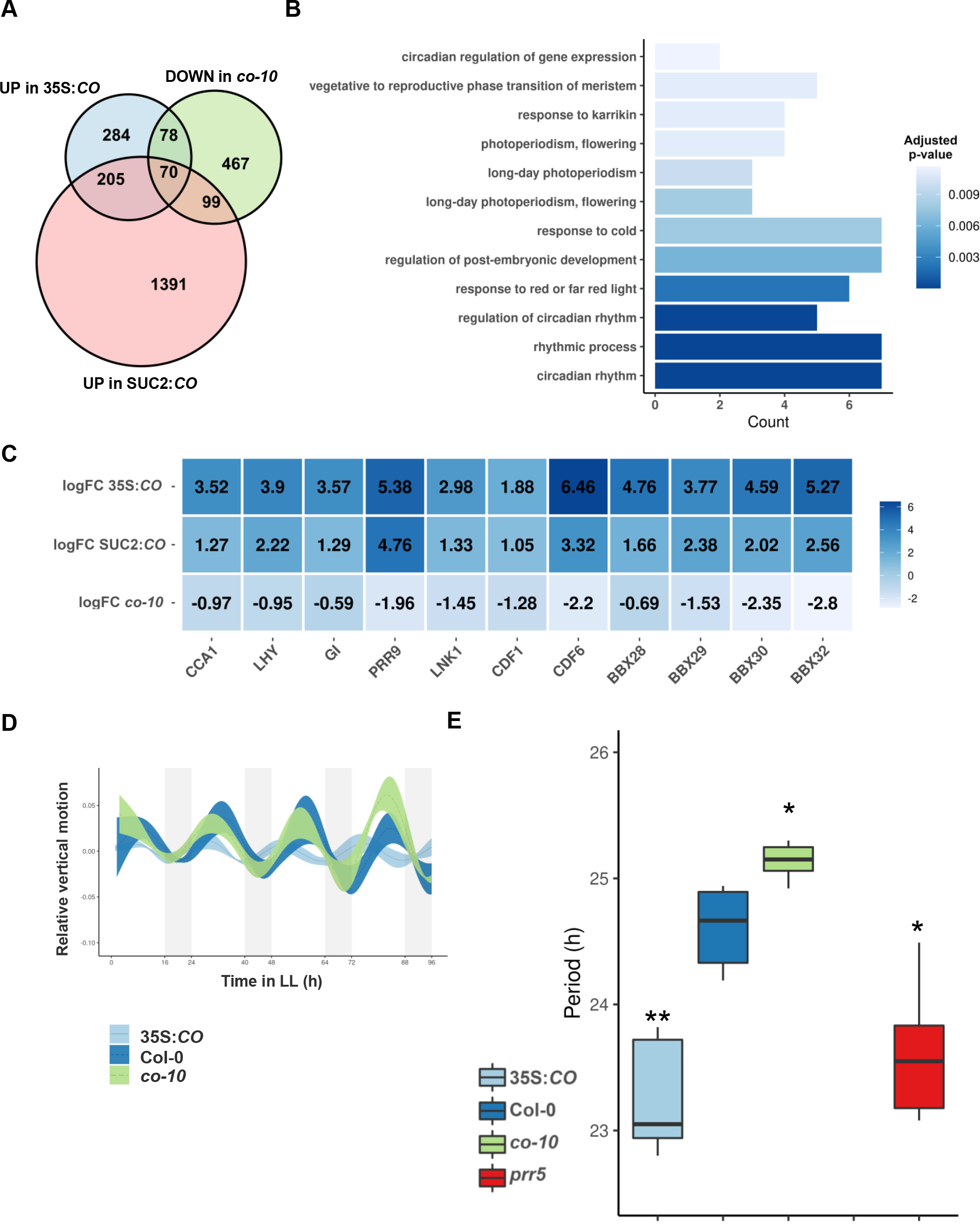
CONSTANS alters the circadian clock system in *Arabidopsis*. (A) Venn diagram representing downregulated genes in the *co-10* mutant (green), upregulated genes in SUC2:*CO* (pink) and upregulated genes in 35S:*CO* plants (blue). The intersecting 70 genes (SuperExact Test, p-value = 1.12 × 10^−114^) are considered high- fidelity CO regulated genes. (B) GO-term enrichment analysis over the intersection between downregulated genes in *co-10* mutants and upregulated genes in CO- overexpressing lines, indicating that they are significantly involved in circadian rhythm processes. (C) Table heatmap showing the change of expression (logFC) between *co- 10*, 35S:*CO*, SUC2:*CO* and the control (WT) for circadian related genes. (D) Relative vertical motion of leaves in 35S:*CO* (light blue), Col-0 (dark blue) and *co-10* (green) plants. Shading indicates the standard deviation. Subjective nights are represented in grey. (E) Period distribution of rhythmic leaf movement in 35S:*CO*, Col-0 and *co-10* plants from the previous image. *prr5 line* is also included as a key circadian clock gene mutant.

To further asses if *CO* was able to modify the circadian pattern of expression of other genes, we analyzed RNA-seq data from a transcriptome of Col-0 plants under LD conditions in a 24 h period (GSE43865, Rugnone et al., 2013) and estimated gene expression levels at ZT2, ZT6, ZT10, ZT14, ZT18 and ZT22. Using this analysis, we identified genes exhibiting a circadian pattern of expression and classified them into clusters depending on the time point at which they reached their minimal and maximal expression (see methods). To identify the specific circadian patterns affected by *CO*, we studied the statistical significance of the intersections between the circadian gene clusters and the activated genes in 35S:*CO* at ZT14. Only three circadian clusters presented a significant intersection according to Fisher’s exact test, as shown in Table S5. All the clusters showing a significant intersection presented a morning peak and an evening/afternoon trough. We concatenated these clusters into a single group called the “morning-phased” cluster and, again, the intersection of these genes with the activated genes in 35S:*CO* was significant (Figure S3A), in agreement with previous studies (Gnesutta et al., 2017). This result shows that CO activates the expression of genes in the evening that naturally exhibit their minimum expression at that time of day, thus altering their circadian patterns.

To confirm the effect of CO on the clock, using RT-qPCR, we measured the expression of some of the genes regulated by CO every 4 h during a 24 h period in WT, *co-10* mutant and 35S:*CO* plants (Figure S3B). We found that *PRR5*, *PRR7*, *GI* and clock-output genes such as *BAM9* were activated in 35S:*CO*, especially at ZT14, when CO protein is stabilized under LD (Suárez-López et al., 2001) and down in *co-10* in *PRR5* and *BAM9*. This reflected the fact that *CO* overexpression, and its absence, altered the daily expression of some clock genes.

To further asses the effect of photoperiod on the clock, we tested if CO was able to alter physiological aspects of the circadian behavior of *Arabidopsis*. In plants, one of the most well-known and characterized circadian clock outputs is the daily rhythmic movement of leaves that has been used to identify several key components of the circadian clock (Wang and Tobin, 1998). We tracked this movement in LD entrained 7- DAG seedlings from WT, *co-10* and 35S:*CO* plants transferred to LL and estimated the period and phase of circadian parameters. As shown in the graphic in Figure 2D and its quantification in Figure 2E, *co-10* mutant showed a longer period than WT, and 35S:*CO* plants exhibited a significant shorter period, similar to *prr5* mutant. The change in nastic rhythms shown in *co* mutants is a strong evidence that CO is involved in circadian clock modification.

To determine if CO could be altering the circadian rhythms by direct binding to clock gene promoters, we further analyzed the ChIP-seq data from 35S:*CO* and *co-10* plants at DAG7. The CO ChIP peak distribution was found to be enriched mainly in promoter regions near the transcription start sites (65%), and to a lesser degree, regions up to 2 kb away (10%), downstream regions (6%) and distant intergenic regions (7%), indicating that it is probably a *bona fide* transcription factor (Figure 3A). Next, we compared the ChIP-seq data with the upregulated genes in 35S:*CO* and SUC2:*CO* (Figure S4A), obtaining a significant overlapping. We found that almost 23% (550 out of 2418) of the CO target genes were also affected in both overexpressor lines, including its main target, the floral integrator *FT*, as well as other photoperiodic growth/development related genes, such as *CDF3* and *CDF6,* and key circadian clock genes, such as *PRR5*, *LHY, GI*, *RVE8* and *RVE4* (Figure S4B). Binding of CO to clock gene promoters suggested that CO was able to alter not only photoperiodic genes expression, as previously suggested (Onouchi et al., 2000), but also the expression of circadian clock-related genes.

**Figure 3.**
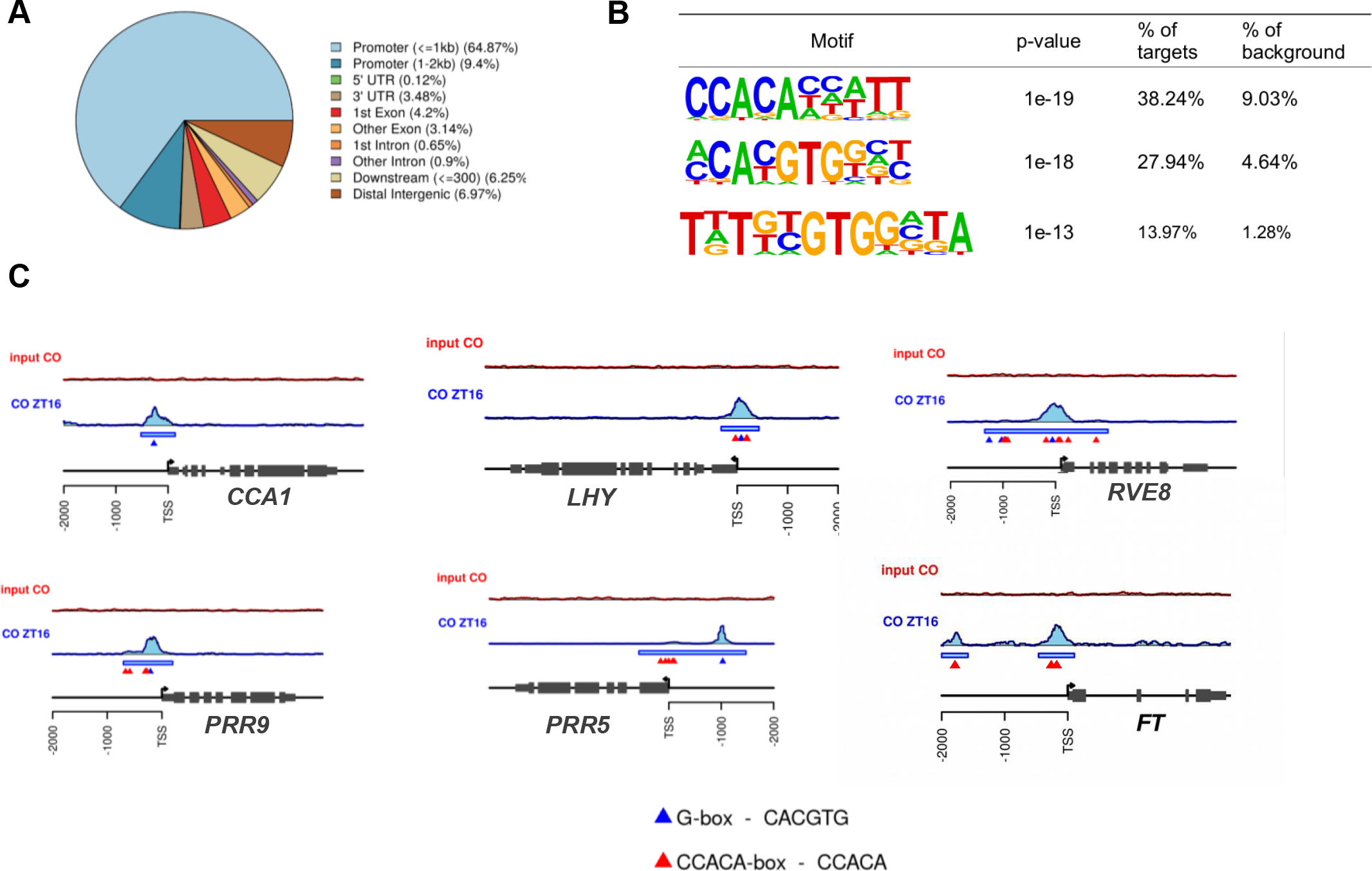
CO binds to clock gene promoters. (A) Pie graph representing the genome-wide distribution of CO peaks. Most of the binding sites (64.87%) are located in promoters (<=1 kb). (B) *De novo* DNA motif identification over the CO binding sites using *HOMER* software and the % of distribution in CO promoters. (C) ChIP-seq visualizer views of CONSTANS occupancy of circadian related genes. The binding profile of immunoprecipitation and input experiments are colored red and blue, respectively. Peaks detected by *MACS* are indicated by blue rectangles. CCACA (red triangles) and G-box elements (blue triangles) located at peaks are also showed.

CO regulates *FT* expression by binding to its promoter in association with NF-Y/HAP heterodimers to bind to CCAAT elements (Wenkel et al., 2006) and to CORE sites (Tiwari et al., 2010). However, recently, it was shown that CO imparts DNA sequence specificity to the NF-Y heterodimer resembling a histone pocket, thus allowing the complex to bind to the CCACA element (Gnesutta et al., 2017, Shen et al., 2020). We performed unbiased DNA motif discovery analysis in the CO ChIP-seq data and confirmed the presence of the CCACA element (p>10^-19^) and another element resembling a CORE-like element (p>10^-13^) (Figure 3B); unexpectedly, we identified a G-box (CACGTG) that was significantly enriched (p>10^-18^) in CO binding peaks. Our analysis showed that CO could bind to G-box sites in *CCA1, LHY* and other loci (Fig 3C) but not on the *FT* promoter, where it was associated exclusively with CCACA sites, revealing a NF-Y/HAP-independent mechanism.

Together, these results suggested that CO is not only an output of the circadian clock pathway but also a factor exerting feedback regulation over key circadian clock genes and circadian patterns, implying a novel function for CO in altering rhythmic processes in *Arabidopsis*.

### CO, PRRs and HY5 share genomic binding sites in circadian clock regulated genes

G-box motifs are the binding sites of bZIP and bHLH TFs and have been described in the transcriptional regulation exerted by PRRs, thus suggesting an association with these TFs (Liu et al., 2016). Under short days (SD, 8 h light, 16 h dark), with a long dark period, the night-inducible bHLH PIFs have been described to bridge the binding of PRRs to DNA to sequentially repress shared target genes related to growth, such as *CDF5* (Martín et al., 2018). Additionally, PIF4 is involved in the temperature-dependent activation of *FT* through binding to CO under SD (Fernández et al., 2016), but, as PIFs are unstable in the light, no involvement on the clock under LD was reported. On the other hand, the bZIP LONG HYPOCOTYL 5 (HY5) is a light-inducible TF (Osterlund et al., 2000) that binds to G-boxes (Lee et al., 2007; Young et al., 2008; Zhang et al., 2011; Binkert et al., 2014), and *hy5* mutations affect flowering (Bhagat et al., 2021) and the clock (Hajdu et al., 2018). It has also been described that HY5 can bind to BBX family proteins from group IV (BBX21-25) through the b-box domain to activate or inhibit its role in photomorphogenesis (Datta et al., 2007; Job et al. 2018; Bursch et al., 2020). Therefore, HY5 was a good candidate to act as bridge mediating the binding of PRR5 and CO to clock-related gene promoters during the day under LDs. HY5 is also degraded by the COP1/SPA1 complex during the night (Osterlund et al., 2000), similar to CO (Jang et al., 2008), and it has been shown that CO and HY5 interact with the diurnal chromatin remodeling factor PKL (Jing et al., 2013; Jing et al., 2019).

Indeed, when we compared the genome-wide analysis results for the ChIP-seq data for PRR5 (Nakamichi et al., 2012) and ChIP-on-chip data for HY5 (Lee et al., 2007) with the ChIP-seq data for CO, a significant overlap between the direct targets of PRR5, HY5 and CO was found (Figure 4A). After carrying out a DNA motif discovery analysis over the promoter sequences (1 kb) of the different gene sets (Figure 4A, right; Table S6), we found a motif resembling the CCACA element (Gnesutta et al., 2017) in the gene promoters regulated exclusively by CO (set 1), indicating that it binds to these regions via an NF-CO trimer. A matrix containing a G-box motif was enriched in the genes regulated by PRR5 (set 2), as described in previous studies (Liu et al., 2016). Notice that the most significant enriched DNA matrices come from bHLH TF ChIP-seq studies (Table S6), suggesting that PRR5 regulates the transcription of these genes in association with bHLHs (Martín et al., 2018). However, a DNA matrix containing an E-box motif ACGTG was recovered from the genes regulated by HY5 (set 3). This suggests a certain degree of specificity imparted by the protein with which HY5 associates. We also found a G-box motif enriched in the gene promoters bound by CO, HY5 and PRR5 (set 123), indicating that this DNA element plays a key role in the transcription of these clock-regulated genes. Furthermore, the most significant DNA matrices were described in bZIP TF ChIP-seq studies (Table S6).

**Figure 4.**
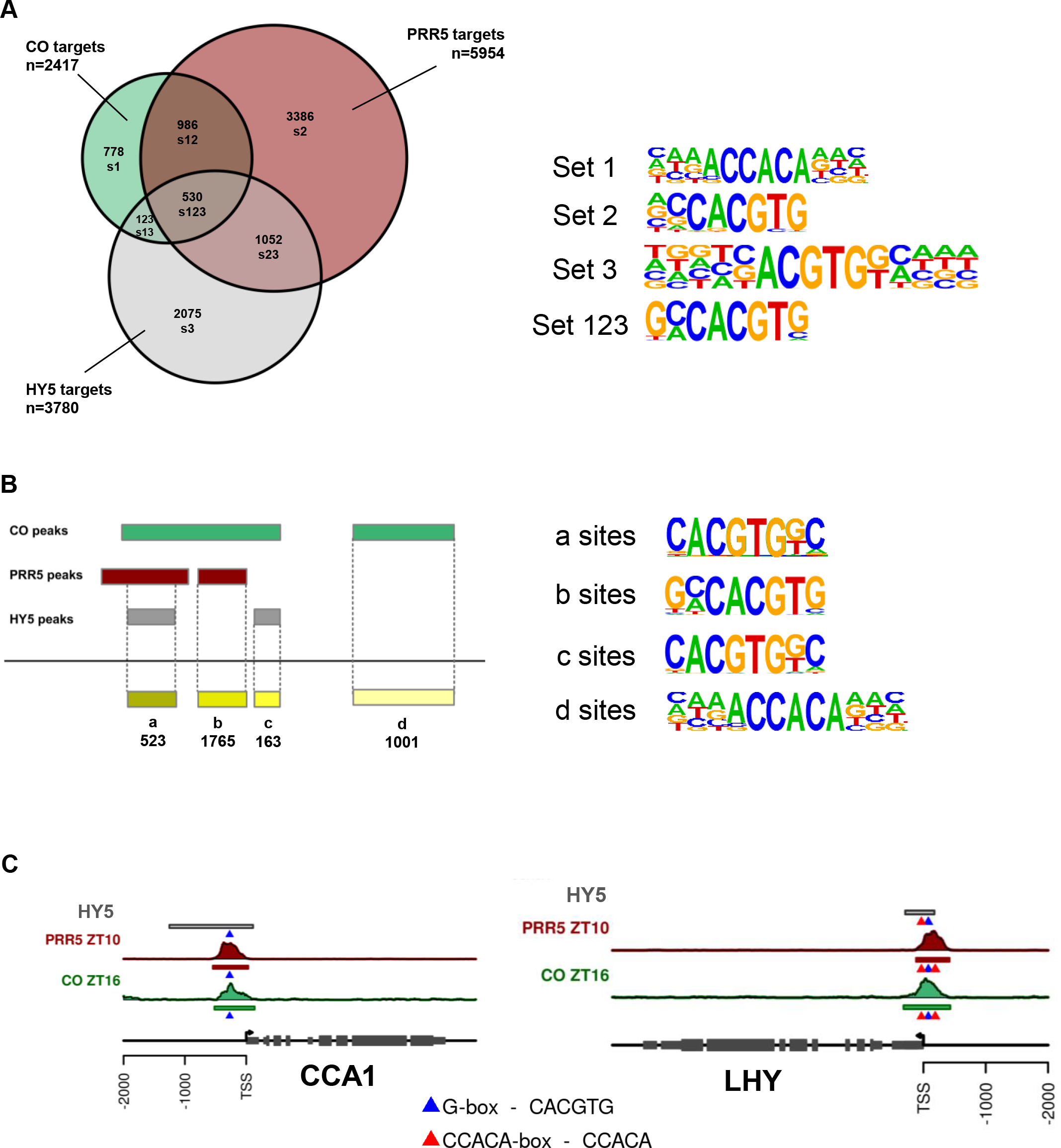
**HY5, PRR5 and CO genome binding analysis**. (A) Left: Venn diagram showing the significant intersection (SuperExact Test, p-value < -10^6^) of CO (green), HY5 (grey) and PRR5 (red) targets. Right: DNA motifs enriched in the promoters of the different gene sets. (B) Left: Graphic scheme of the peak intersection analysis. Right: DNA motif enrichment analysis performed in each retrieved peak set. (C) Peak visualizer views of HY5 (grey), PRR5 (red) and CO (green) binding to *CCA1* and *LHY* promoters. G-box and CCACA elements are indicated by blue and red triangles.

To support our hypothesis, we followed a similar approach to analyze the genome-wide binding sites of CO, HY5 and PRR5 (Huang et al., 2012; Nakamichi et al., 2012; Liu et al., 2013; Liu et al., 2016) (Table S7). First, we found a significant overlap between the three sets of binding regions according to permutation tests (Table S8). Next, using the *intersectBed* function from *bedtools*, we retrieved the overlapping peaks for the three TFs (a), the intersection between CO and PRR5 sites (b), the intersection between HY5 and PRR5 sites (c) and the genome regions bound by CO (d) (Figure 4B, left). In order to determine the DNA elements involved in the regulation exerted by these TFs, we performed DNA motif discovery analyses over the different binding site sets (Figure 4B, right). A CCACA element (Gnesutta et al., 2017) was enriched exclusively in the CO binding sites (d), which is consistent with previous studies based on transcriptomic data (Gnesutta et al., 2017). However, G-boxes were enriched in the other intersections, a-c sites (Figure 4B, right). Furthermore, we analyzed ChIP-seq data for other PRR family members at different time points. We found an overlap between the binding sites of CO, HY5, TOC1, PRR5, PRR7 and PRR9 in G-box-containing sites located in clock promoters (Figure S5A). All these results indicate that CO and PRR5 bind to a G-box to control the transcription of clock-regulated genes, probably in association with a bZIP TF, such as HY5, as shown in Figure 4C for the *CCA1* and *LHY* promoters.

To further demonstrate HY5 binding to these sites, we performed ChIP experiments using GFP-Trap antibodies in 35S:*GFP*:*HY5* (*hy5-215*) plants and observed a significant enrichment of these amplicons compared with the control (WT plants) that was absent in the *FT* promoter (Figure S5B). These results indicated that HY5 binds to the same sites as CO in these circadian clock promoters but not the CORE site associated with CO in the promoter of *FT*, highlighting that CO has a different function related to circadian rhythm that is independent of the photoperiodic flowering pathway.

To further confirm the role of HY5 in CO-dependent clock gene transcriptional modification, we crossed the *hy5-2* mutant with 35S:*CO* plants to obtain the double 35S:*CO hy5* mutant (Figure 5A). These plants showed a small delay in flowering time compared to 35S:*CO* plants (Figure 5B). 35S:*CO hy5* plants showed high expression of *CO* and no expression of *HY5* mRNA or protein, as confirmed by RT-qPCR (Figure S6A) and Western blot experiments (Figure S6B), respectively. Consequently, the expression of the clock-related genes *CCA1*, *PRR5, PRR7* and *GI* activated by CO was reduced (Figure 5C), further confirming the role of HY5 in CO-dependent clock gene expression.

**Figure 5.**
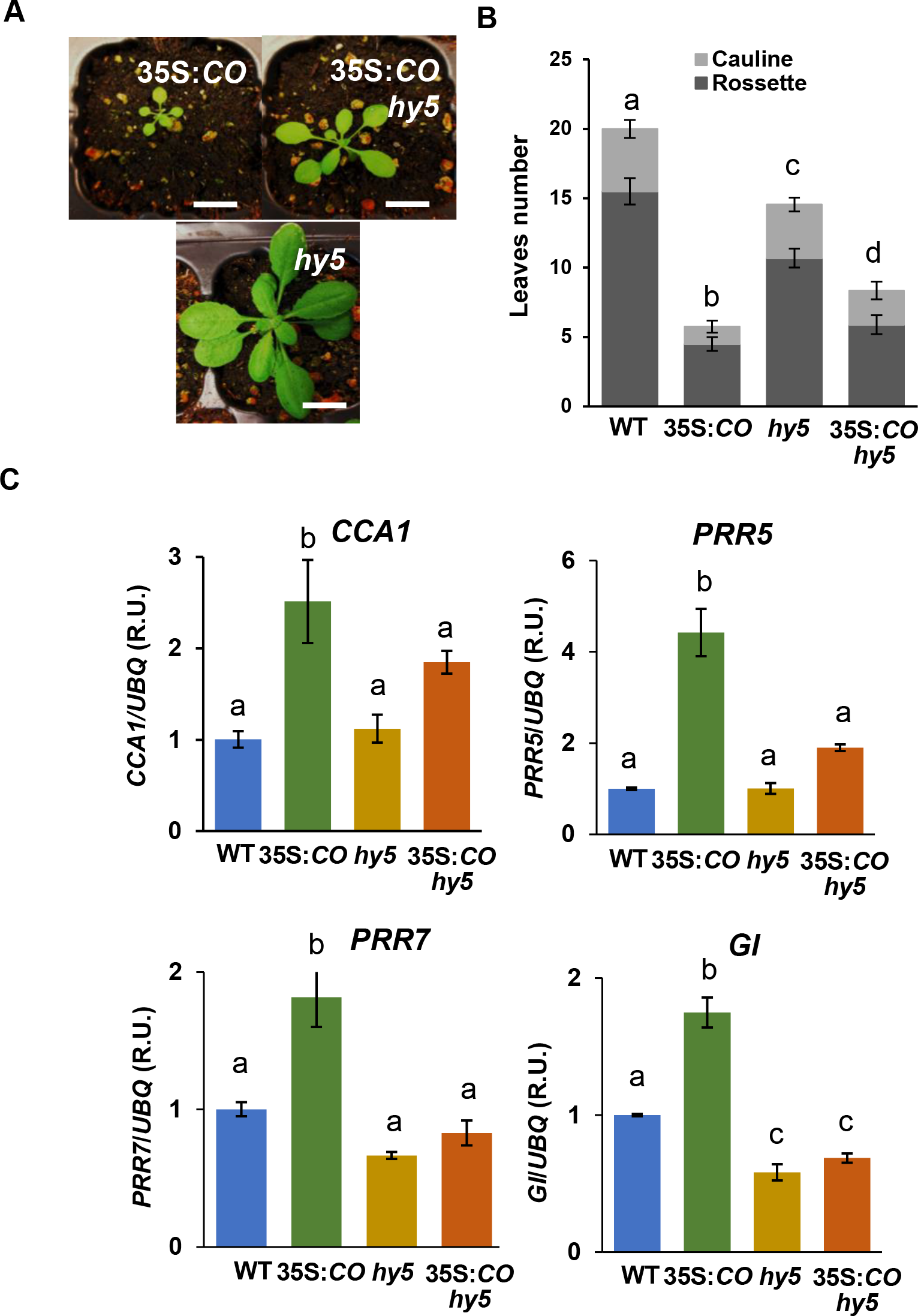
Genetic interaction between *CO* and *HY5*. (A) Pictures of 35S:*CO*, *hy5-2* and 35S:*CO hy5* plants at DAG15. (B) Flowering time of WT, 35S:*CO*, *hy5-2* and 35S:*CO hy5* plants measured as total leaf number. (C) Expression of *CO* target genes *CCA1, PRR5, PRR7* and *GI* in WT, 35S:*CO*, *hy5-2* and 35S:*CO hy5* plants at ZT14, DAG 15. Error bars indicate the standard deviation (s.d.) from three independent experiments. P < 0.05, one-way ANOVA and Tukey’s HSD.

### CO, PRR5 and HY5 proteins interact *in planta*

The interaction between CO and some members of the PRR family was previously shown; this binding stabilizes CO and enhances photoperiodic flowering under LD conditions (Hayama et al., 2017). However, considering that CO and PRR5 bind to the same sites in some promoters and that these two proteins seem to have the opposite effect over transcription, it is possible that CO can gradually remove PRR5 from the DNA and promote transcription, together with HY5. Therefore, a transitory complex consisting of HY5, PRR5 and CO during the evening could exist to alter gene expression. To examine this possible ternary complex, we performed different protein-protein interaction approaches.

First, we performed a bimolecular complementation (BiFC) assay in *Nicotiana* between CO, HY5 and PRR5 in pairs, and we observed the interactions between them in the nucleus and found different distribution patterns (Figure 6A). We also performed yeast-two-hybrid experiments with the three main domains of CO (Bboxes, middle domain and CCT domain) to confirm the interaction with both HY5 and PRR5. As can be seen in Figure 6B, and different from the previous results on the binding of other BBXs to HY5 through the Bboxes (Bursch et al., 2020), CO interacts with HY5 and PRR5 through the middle domain and, to a lesser extent, the CCT domain. To further investigate if there could be a ternary complex, we tested the colocalization of the three proteins in the nucleus of *Nicotiana* cells. To do this, we reconstituted the YFP fused to HY5 and PRR5, and co-infiltrated a *CO*:*CFP* construct. We detected a significantly higher colocalization in speckles, compared with the control with CFP alone (Figure 6C). Following this approach, we measured fluorescent resonance (FRET) between CFP and YFP through a sensitized emission method, and we found higher values in the CO/HY5/PRR5 combination than in the negative controls, confirming the close interaction among the three proteins (Figure 6D).

**Figure 6.**
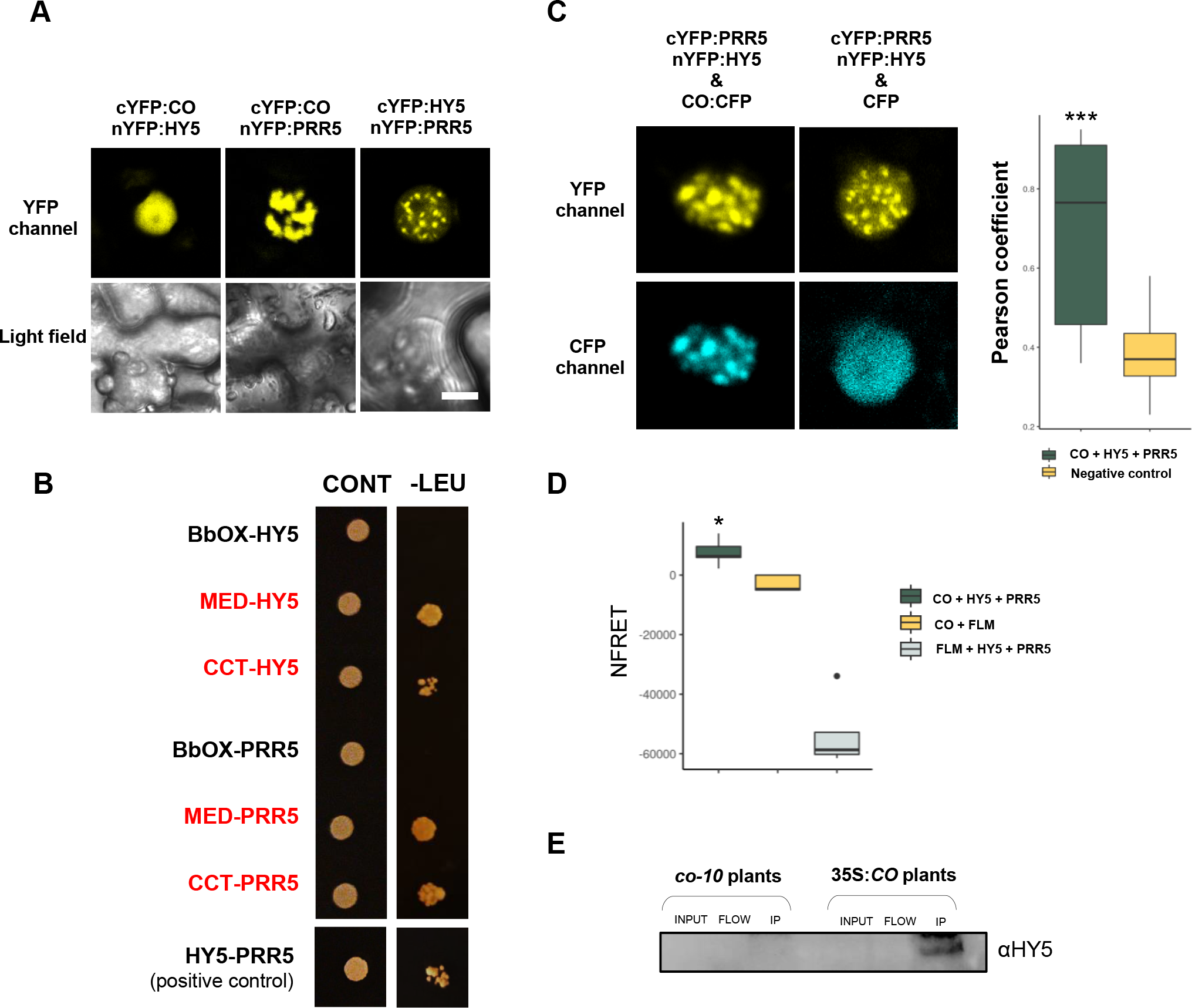
Protein interaction between CO, HY5 and PRR5. (A) Confocal images of CO, HY5 and PRR5 interactions using BiFC assays in *Nicotiana* leaves. (B) Y2H assays showing the interaction between different parts of CO (Bbox, middle and CCT domains) with HY5 and PRR5. Panels show 3-day-old colonies grown in control and selective media. (C) Colocalization experiments of CO, HY5 and PRR5 proteins. YFP was reconstituted by fusing N- and C- terminal parts to HY5 and PRR5; the full-length sequence was fused to CO. The colocalization between both fluorophores was measured by Pearson correlation compared to a negative control. (D) FRET analysis between CO, HY5 and PRR5. CO was fused to CFP and YFP was reconstituted by YFN:HY5/YFC:PRR5. CFP:FLM and YFP:FLM were used as negative controls. The proteins were coexpressed in *N. benthamiana* leaves, and normalized FRET (nFRET) was measured in 10 cells. (E) Immunodetection of the HY5 protein with αHy5 in chromatin immunoprecipitation experiments (using αCO) of 35S:*CO* and *co-10* plants.

Since CO and HY5 physically interact and both proteins bind to the same regions in clock gene promoters, we investigated if they could associate with DNA when combined. To examine this possibility, the immunoprecipitated complex resulting from the ChIP experiment was analyzed by immunoblot using anti-HY5 antibodies. We detected HY5 protein in the 35S:*CO* sample but not in the control (Figure 6E), confirming the presence of a complex between CO-HY5 and DNA.

The clock-related leaf movement was also affected in *PRR5* and *HY5* mutants, showing an acceleration of the period compared to 35S:*CO* plants (Figure S7).

Together, these results indicated that these three proteins could form a complex that binds to the DNA, acting HY5 as scaffold, since this type of TF, bZIP, is largely described to be capable to directly binding to DNA (Chattopadhyay et al., 1998; Abbas et al., 2014; Binkert et al., 2014).

## Discussion

A network science perspective is essential for understanding complex interconnected systems (Barabási, 2015). Transcriptional programs governing circadian clock and photoperiodic pathways showed a high relatedness in CircadianFloralNet according to its scale-free and small-world properties, which revealed many unexplored connections. In the field of graph theory, network motifs are particularly interesting because they confer particular functions. Network motifs are subgraphs that appear in an observed network significantly more frequently than they do in other comparable networks (Milo et al., 2002; Shen-Orr et al., 2002; Stone et al., 2019). Despite the high complexity of gene networks, a certain level of simplicity can be found in these sub- structures, which are present throughout biological networks and across species (Alon, 2007) Much debate has focused on the validity of analyzing these network motifs because they are components of large and complex networks (Mellis and Raj, 2015) Indeed, CircadianFloralNet represents all potential interactions under many conditions, tissues, growth stages and times of day; furthermore, smaller and insulated networks have to be considered. Thus, the network motif formed by *CO* and *PRR5* functions in the evening under LD conditions during the floral transition.

In recent years, the circadian clock in *Arabidopsis* has emerged as a complex regulatory system; it is not a simple molecular timer but a regulator of multiple processes involving hormone, metabolism and stress pathways, some of which exert feedback regulation on the clock (Sanchez and Kay, 2016). Indeed, several TFs that contribute to clock function have been identified (McClung, 2014) and it has been proposed that different tissues may show differences in the clock (Endo et al., 2014).

CO function is strongly regulated at both the transcriptional and post-translational levels in *Arabidopsis*, indicating that the photoperiodic pathway has been subjected to high selective pressure (Romero and Valverde, 2009). Other than flowering transition and reproductive development, no other functions had been described for CO, but recent studies suggested that it could regulate floral jasmonate signaling (Serrano-Bueno et al., 2022), seed size (Yu et al., 2023) and circadian genes (Gnesutta et al., 2017). Here, we show that CO alters the circadian clock by binding to the promoters of key clock-related genes, revealing an undescribed function. Therefore, there is an even more complex regulation between the circadian clock and photoperiod pathways than previously shown.

PRRs have been described as repressors of their direct target genes. According to our network based on ChIP-seq data, PRR5 could bind to the *CO* promoter. Nevertheless, PRRs cause a global positive effect over the photoperiod pathway and CO-FT module regulation, since they stabilize CO (Hayama et al., 2017) and repress *CDF1* expression, triggering the net activation of *CO* and *FT* expression (Nakamichi et al., 2007). Then, the control of *PRR5* gene expression by CO establishes a positive feedback loop between the circadian clock and the photoperiodic pathway. In addition to PRRs, CO binds to the *GI* promoter. Since GI exerts an activating role over CO (Imaizumi et al., 2005; Sawa et al., 2007; Song et al., 2014), this could also form a positive feedback loop to enhance and fix the photoperiodic response in the afternoon under LDs. The crosstalk between flowering and the circadian clock exerted by CO could help to explain the existence of tissue-specific clocks (Endo et al., 2014). These results support the already established concept of complex interlocking loops instead of a unidirectional system composed of input-oscillator-output.

Moreover, it seems that in addition to affecting clock genes such as *PRR5* or *GI*, CO could regulate circadian clock target genes, including a significant number of PRR5 targets. This set of genes would be repressed by PRR5 and activated by CO, forming a transcriptional switch. In order to move from a vegetative phase to a reproductive phase, massive changes in gene expression are needed in plants; thus, CO could provide the activation of a set of repressed genes and promote the phase transition (Figure 7).

**Figure 7.**
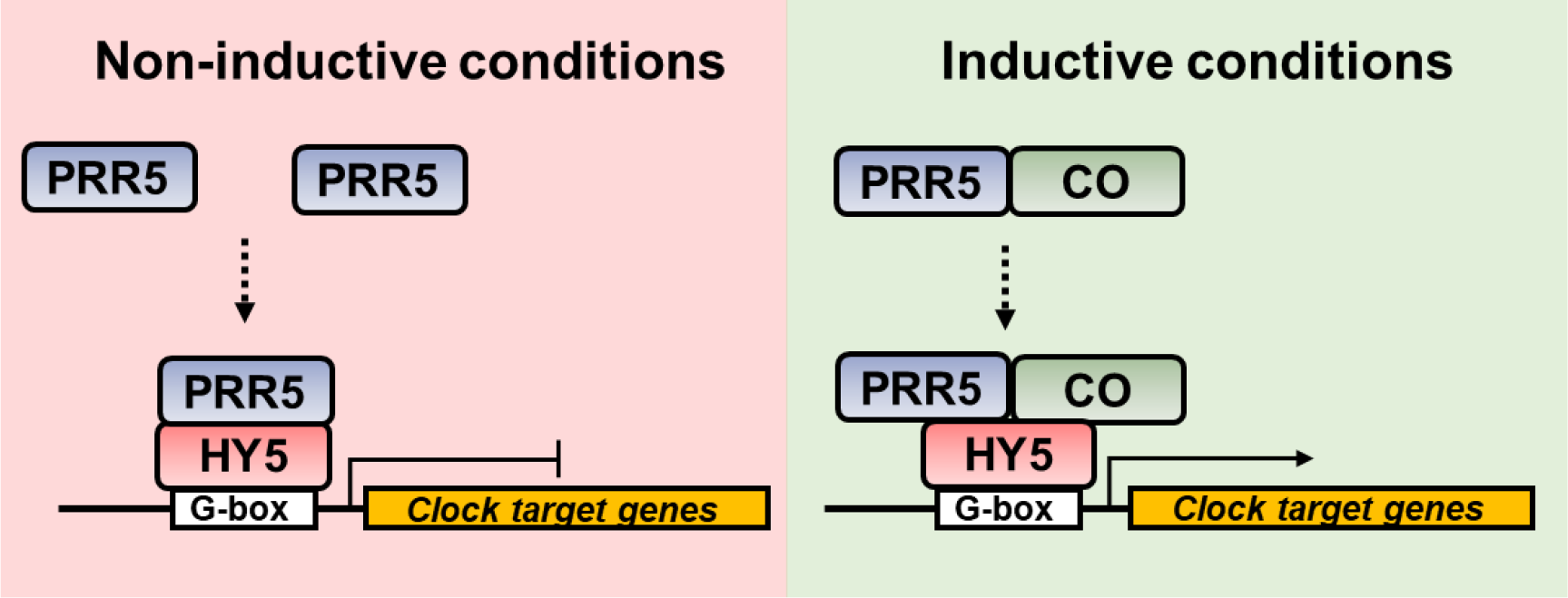
Proposed model of the regulation of the circadian clock by CO. Under noninductive conditions, PRR5 binds to G-boxes in the promoter of its target genes and represses their expression. When CO accumulates (i.e. LD), it binds to PRR5 to stabilize it and occupies the same genome binding sites to activate the expression of its target genes. HY5 or another bZIP transcription factor could mediate CO and PRR5 binding to the DNA.

There has been a long controversy about whether CO could act as a TF or as a coactivator (Blackman and Michaels, 2010). Recently, it has been discovered that CO can directly bind to CORE elements and associate with NF-Y/HAP heterodimers to bind to CCACA elements, acting as an activator of gene expression (Wenkel et al., 2006; Tiwari et al., 2010; Gnesutta et al., 2017). Indeed, CO activity can be controlled by the formation of different types of protein complexes, even repressor complexes formed by TOPLESS (Graeff et al., 2016). Here, we demonstrate that CO is able to bind to G-box elements in DNA through the bZIP transcription factor HY5, a TF involved in many light processes and pathways (Gangappa and Botto, 2016). Together, these findings show a great plasticity in CO function as a transcriptional regulator and its ability to recruit circadian clock machinery at the end of a long day. Depending on its partners, CO will exhibit different effects over transcription and affinity for different genome binding sites; thus, conferring high plasticity to plants in response to daylength changes.

## Methods

### Plant material and growth conditions

Plant lines in the Col-0 background used in this study included 35S:*CO* (Onouchi et al., 2000), SUC2:*CO* (Hayama et al., 2017), *co-10* (SAIL_24_H04), *hy5-2* (SALK_056405C), 35S:*GFP:HY5* (*hy5-211*) (kindly provided by Dr Vicente Rubio), and *prr5-1* (SALK_006280; Michael et al., 2003). 35S:*CO hy5* double mutant plants were generated by crossing 35S:*CO* with *hy5-2* to form homozygotes. *Arabidopsis* seedlings were grown in a SG-1400 phytotron (Radiber SA, Spain) under a LD light regime with temperatures ranging from 22°C (day) to 18°C (night) and 100 μEm^-2^ s^-1^ light intensity.

*Arabidopsis* seeds were stratified for 3 days at 4°C in the dark and later grown in MS medium. For gene expression analyses, *Arabidopsis* Col-0, 35S:*CO* and *co-10* seedlings were collected every 4 h during a complete LD. For monitoring leaf movement, seedlings were grown in soil for 10 days under LD conditions, transferred to LL conditions and examined for 4 days.

### Chromatin immunoprecipitation experiments

ChIP was carried out as described previously (Bowler et al., 2004) with minor modifications to improve binding enrichment (Lau and Bergmann, 2015). Plants were grown for 7 days under LD conditions and 9 g of seedlings were harvested at ZT12- ZT16. Then, plant tissue was crosslinked in a vacuum concentrator in presence of 1 mM DSG for 10 min, 1% (v/v) formaldehyde for 20 min and finally 0.125 M glycine to stop crosslinking. Later, the tissue was ground in liquid nitrogen and chromatin was isolated. Chromatin complex immunoprecipitation was performed using αCO (Serrano-Bueno et al., 2020) and GFP-Trap agarose beads (ChromoTek) to capture CO and HY5 proteins, respectively. After immunocomplex capture, DNA was eluted from the beads, crosslinked with proteins, reverse transcribed, and isolated. The subsequent qPCR was performed as previously described. Immunoprecipitated samples were normalized to a 10% reverse crosslinked fraction of each chromatin preparation. Primers used to measure DNA fragments enriched during ChIP experiments are shown in Table S9. Two biological replicates were processed for next-generation library preparation using the ThruPLEX DNA-seq kit (Takara). Next, libraries were sequenced on NextSeq500 (Illumina) at the Cabimer CSIC Genomics Unit. CO ChIP-seq analysis was implemented as described below. The raw data has been deposited in Gene Expression Omnibus (GEO) under accession GSE222657.

### ChIP-seq data acquisition and analysis

We used ChIP-seq data generated for transcription factors involved in flowering and the circadian clock that were available from the Sequence Read Archive (SRA, https://www.ncbi.nlm.nih.gov/sra) (Wheeler et al., 2005). Specifically, data from 33 different transcription factors were downloaded in this study. These transcription factors and their ChIP-seq data accession numbers are shown in Table S10. The *Arabidopsis* reference genome TAIR10 and its corresponding annotation were downloaded from the Ensembl Plants website (http://plants.ensembl.org/index.html). The FASTQC software package was employed to examine the read quality of each sample (http://www.bioinformatics.babraham.ac.uk/projects/fastqc/). The *fastq* files corresponding to each transcription factor were mapped to the reference genome using the ultrafast and memory-efficient short read mapper *bowtie* (Langmead et al., 2009). The read mappings for each sample were stored in *SAM* (Sequence Alignment Map), *BAM* (Binary Alignment Map) and *bigWig* files. We performed the “peak calling” step with the software tool *MACS2* (Zhang et al., 2008). Finally, target gene selection was carried out using *PeakAnnotator*, part of the *PeakAnalyzer* program (Salmon-Divon et al., 2010), and an *ad-hoc* R script developed in this study according to the criterion of the nearest downstream gene. DNA motif identification was carried out using *HOMER* software (Heinz et al., 2010). For further peak annotation and visualization, *ChIPseeker* (Yu et al., 2015) and *ChIPpeakAnno* (Zhu et al., 2010) R packages were employed. Finally, GO term enrichment analysis was performed using the R package *clusterProfiler* (Wu et al., 2021). For the CO ChIP-seq analysis, peaks detected in the 35S:*CO* experiment were filtered based on the peaks found in *co-10* immunoprecipitation.

### Transcriptional network construction and analysis

Our transcriptional network was generated and analyzed with the R package *igraph* (Csardi, 2006). Graphical representations of the network were done using *Cytoscape* (Smoot et al., 2011), a software package for network visualization and data integration. Specifically, the ‘prefuse force layout’ method was applied for network visualization. Additionally, a network topology analysis was achieved. The network node-degree distribution was calculated through degree function from the *igraph* package. To analyze if the network adjusts to the scale-free property, the power.law.fit function (*igraph* package) was used to test the node degree distribution and determine whether it followed a power law. Furthermore, the network clustering coefficient and the average path length was calculated through transitivity and average.path.length functions, respectively. A network motif is a subgraph that appears a significantly higher number of times in the network of interest when compared to similar random networks. Therefore, we tested the significance of all possible subgraphs consisting of one, two or three nodes by generating 100,000 random graphs with the same topological features as CircadianFloralTFNet.

### RNA isolation and gene expression analysis

Total RNA from *Arabidopsis* seedlings (0.1 g leaf tissue) was isolated by the Trizol (Invitrogen) procedure following the recommendations of the manufacturer. The final RNA sample was suspended in 21 µL of diethyl pyrocarbonate-treated water, quantified in an ND-1000 spectrophotometer (Nanodrop) and stored at −80°C. Then, 500 ng of RNA was used to synthesize cDNA employing the QuantiTect Reverse Kit (Qiagen) and stored at −20°C until qPCR was performed. qPCR was performed with the iQTM5 Multicolor Real-Time PCR Detection System (Bio-Rad) in 10-µL reactions: primer concentration of 0.2 µM, 10 ng of cDNA, and 5-µL of SensiFAST SYBR & Fluorescein Kit (Bioline). Each sample was measured in triplicate. The qPCR program consisted of (1) one cycle of 95°C for 2 min; (2) 40 cycles of 95°C for 5 s, 60°C for 10 s, and 72°C for 6 s; and (3) one cycle of 72°C for 6 s. Fluorescence was measured at the end of each extension step, and melting curve analysis was performed between 55°C and 95°C. The initial concentrations were calculated using the ddCt Algorithm (Zhang et al., 2016). Primers used to measure gene expression by qPCR are listed in Table S9.

### RNA-seq data acquisition and processing

In this study, we generated RNA-seq data and used other publicly available at the Sequence Read Archive (SRA). We analyzed different studies (GSE43865, GSE236178 and GSE205675) and obtained 31 samples representing three different genotypes, Col-0, *co-10*, SUC2:*CO* and 35S:*CO,* under LD and LL conditions (see supplementary material: Table S11). The quality control of each sample was carried out using the FASTQC software, and the read sequences were mapped to the reference genome with the ultrafast universal aligner *STAR* (Dobin et al., 2013). The number of reads that mapped to each gene was perform using the *featureCounts* software of the *Subread* package (Liao et al., 2014). Genes with less than 10 reads in all samples were removed, and TMM normalization and log2 transformation were performed with *calcNormFactors* (*edgeR* package; Robinson et al. 2010) and *voom* functions (*limma* package; Ritchie et al., 2015), respectively. Next, a linear model with the variable ‘genotype’ was fitted for each gene using the *lmFit* function (*limma* package) in order to carrying out the differential expression analyses. The differentially expressed genes (DEGs) were identified using a threshold of 1.5-fold change and an adjusted p-value < 0.05.

### Protein analysis

Proteins were isolated from seedlings grown on MS-agar using the TRIZOL (Invitrogen) protocol, separated by SDS-PAGE by standard procedures, transferred to nitrocellulose membranes, incubated with αCO (Serrano-Bueno et al., 2020) or αHY5 (Agrisera) antibodies, and developed with a chemiluminescent substrate (Clarity^tm^ Western ECL, Bio-Rad). Blots were visualized and quantified in an Amersham Imager 680.

### Determination of circadian genes and clustering techniques

The *bioconductor* R package *RAIN* (Rhythmicity Analysis Incorporating Non- parametric methods) (Thaben and Westermark, 2014) was used to determine circadian genes, namely genes exhibiting rhythmic patterns of expression with a period of 24 h. A time interval of 4 h and a p-value of 0.01 were used. Circadian genes were classified into clusters based on the positions of their peak (maximal level of expression) and trough (minimal level of expression) during a 24 h period.

### Circadian leaf movement analysis

After transfer to LL, the first pair of leaves was monitored using a Raspberry Pi camera. The leaf movement was tracked using *TRIP* software (Greenham et al., 2015) run in Octave, and the vertical motion was obtained as a function of time and analyzed using *BioDare* software (which performs FFT-NLLS period analysis) (Zielinski et al., 2014) and the *CircaCompare* package (Parsons et al., 2020) implemented in R.

### Transitory expression in *Nicotiana benthamiana*

To perform BiFC assays, *CO, HY5, PRR5* and *FLM* cDNAs were cloned into pYFN43 and pYFC43 Gateway vectors (Belda-Palazón et al., 2012) producing a N-terminal fusion with nYFP and cYFP protein, respectively. *A. tumefaciens* strain GVG3101 pmp90 were transformed with these constructs, and 4-week old *N. benthamiana* plants were infiltrated. Later, leaf disks were visualized with the Leica TCS SP2/DMRE confocal microscope. YFP was excited with the 514 nm line of an argon laser at 20% strength, and fluorescence was detected between 515-565 nm. For ternary complex colocalization and FRET experiments, *HY5* and *PRR5* cDNAs were fused to C- and N- terminal fragments of YFP as in the BiFC assay, and *CO* and *FLM* cDNAs were cloned into pGWB644 and pGWB642 vectors, which produces a C-terminal fusion of full- length CFP and YFP, respectively (Shyu et al., 2008). CFP was excited with the 458 line under a Leica TCS SP2 confocal microscope, and the emission was detected between 465-479 nm. YFP was excited with a 514 nm laser, and the band-pass filters were adjusted to 520-545 nm. Colocalization was measured using the Pearson coefficient of correlation implemented in the *coloc2* plugin of the Fiji distribution of *ImageJ* (Schindelin et al., 2012).

### Y2H assays

The DupLEX system was used to detect protein-protein interactions between CO domains and PRR5 or HY5. CO domains (CCT, middle and Bbox domains) were cloned into the bait vector pJG4-5, while full-length CDSs of *PRR5* or *HY5* were cloned into the prey vector pEG202. Primers used to generate Y2H clones are listed in Table S9. EGY48 cells (MATα *trp1 ura3 his3 LEU2*::pLex *Aop6*-*LEU2*) were used as the host strain for Y2H experiments (Gyuris et al., 1993). Single colonies grown on selection plates were inoculated in 3 ml of SD-Ura-His-Trp and grown overnight at 28°C. Saturated culture was then used to make serial dilutions with an optical density at 600 nm (OD600) of 4^-1^. Then, 2.5 µl of the solution was then spotted on an SD-Ura-His-Trp plate, with an SGalRaf-Ura-His-Trp-Leu plate as a growth control. Plates were imaged after incubation for 60-72 h at 30°C.

### Statistical analysis

The mean comparison between two sets of data was performed using the Student’s t-test after test the normality of the data with the Shapiro-Wilk test. The difference between means was considered statistically significant when the p-value was less than 0.05 (marked with a single asterisk), 0.01 (marked with two asterisks) or 0.001 (marked with three asterisks). Fisher’s exact test was used to analyze the significance of Venn diagram overlaps. Precisely, shapiro.test, t.test and fisher.test functions from R package stats were employed. To determine the significance of overlapping peaks, Monte Carlo permutation tests were carried out, in which one of the peak sets was randomly shuffled and new intersections were calculated after each shuffle; 10000 iterations were computed in order to estimate p-jvalues. These p values were corrected for multiple testing using the Benjamini-Hochberg method. Statistically significant differences between means were evaluated using ANOVA R package.

## Acknowledgements

Work by G. S.-B. was supported by a European Union contract LONGFLOW, MSCAIF- 2018- 838317 and CSIC LONGFLOW, CONV_EXT_014. The financial support of the Spanish Ministry for Science and Innovations (MICINN/FEDER) PID2020-117018RB- I00 to F.V. is also acknowledged. We thank Prof. George Coupland (MPIPZ, Cologne, Germany) for discussion and critical reading of the manuscript.

The authors declare no competing interests.

---

**Figure S1.**
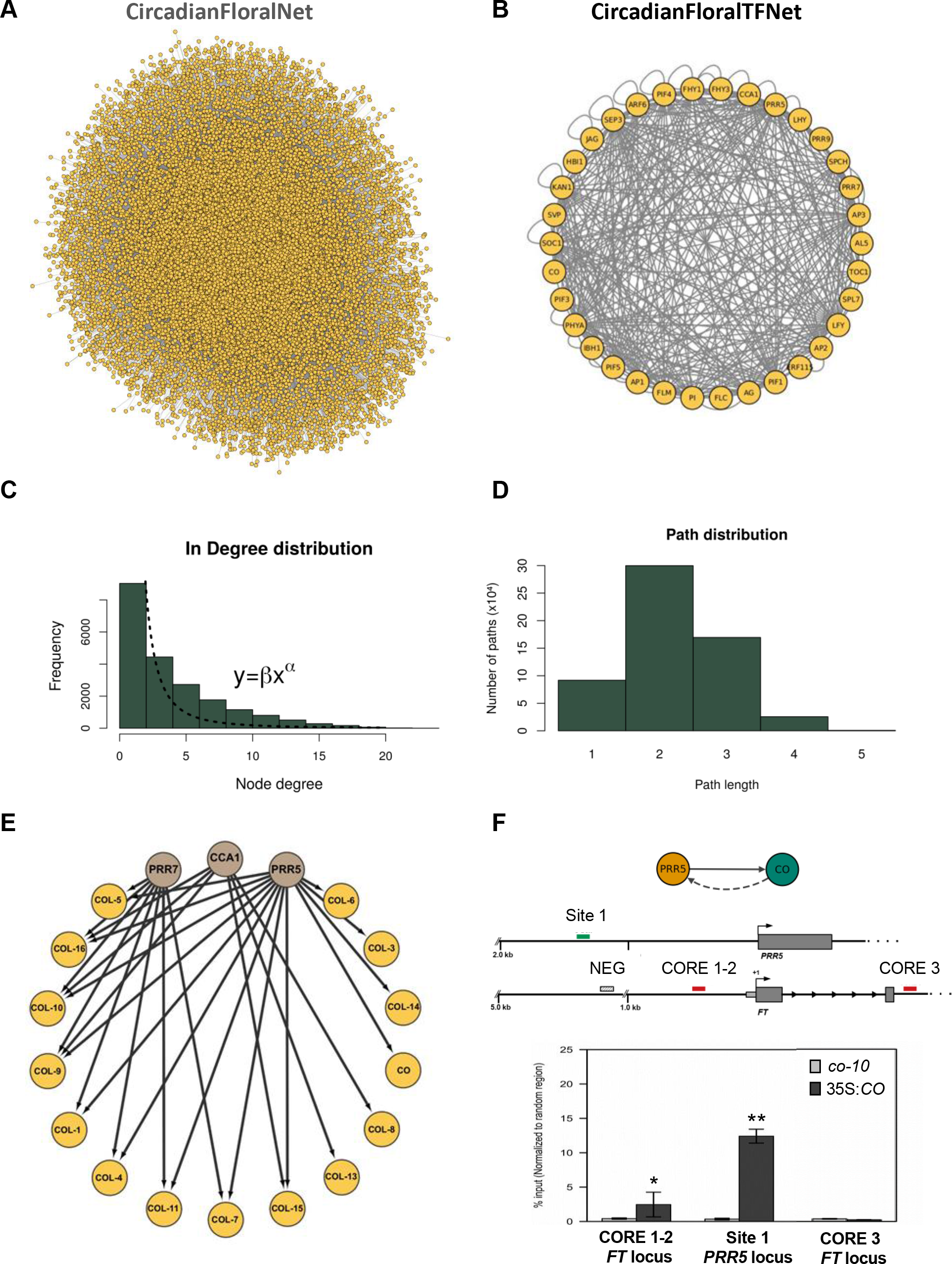
Transcriptional network analysis. (A) *Cytoscape* representation of CircadianFloralNet, a transcriptional network integrating ChIP-seq data from TFs involved in the circadian clock, flowering transition and floral development. (B) CircadianFloralTFNet, the regulator core composed only of the analyzed TFs and their interactions. (C) In-Degree distribution of CircadianFloralNet, following a power law. (D) Path distribution of CircadianFloralNet showing that the average path distance between two random nodes is 2.089. (E) Subnetwork representing the transcriptional interactions between the *CO-like* gene family and central circadian regulators (*CCA1*, *PRR7* and *PRR5*). (F) ChIP-QPCR (below) of amplicons located in the *FT* and *PRR5* promoters (above) with the CO protein.

**Figure S2.**
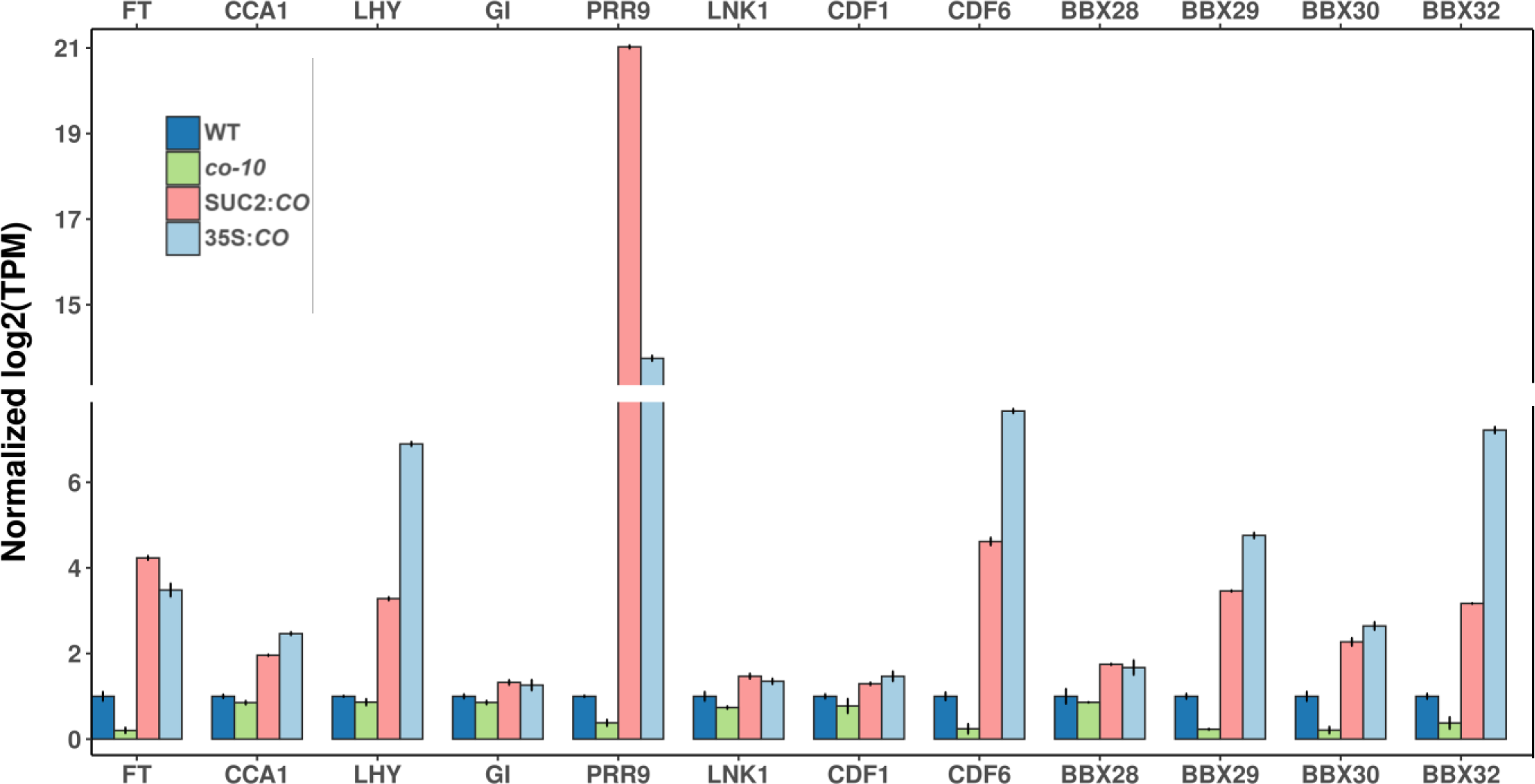
Normalized expression level in Col-0 (dark blue), *co-10* (green), SUC2:*CO* (pink) and 35S:*CO* (light blue) based on the RNA-seq experiments for circadian clock- and photoperiod-related genes. Since these datasets come from two different experiments (Col-0 and *co-10* from GSE205675; Col-0, SUC2:*CO* and 35S:*CO* from GSE236178), the value of Col-0 was normalized to 1 for each gene. *PRR5*, *PRR7*, *GIGANTEA* and *BAM9* expression levels during a LD time course in Col-0, 35S:*CO* and *co-10* lines.

**Figure S3.**
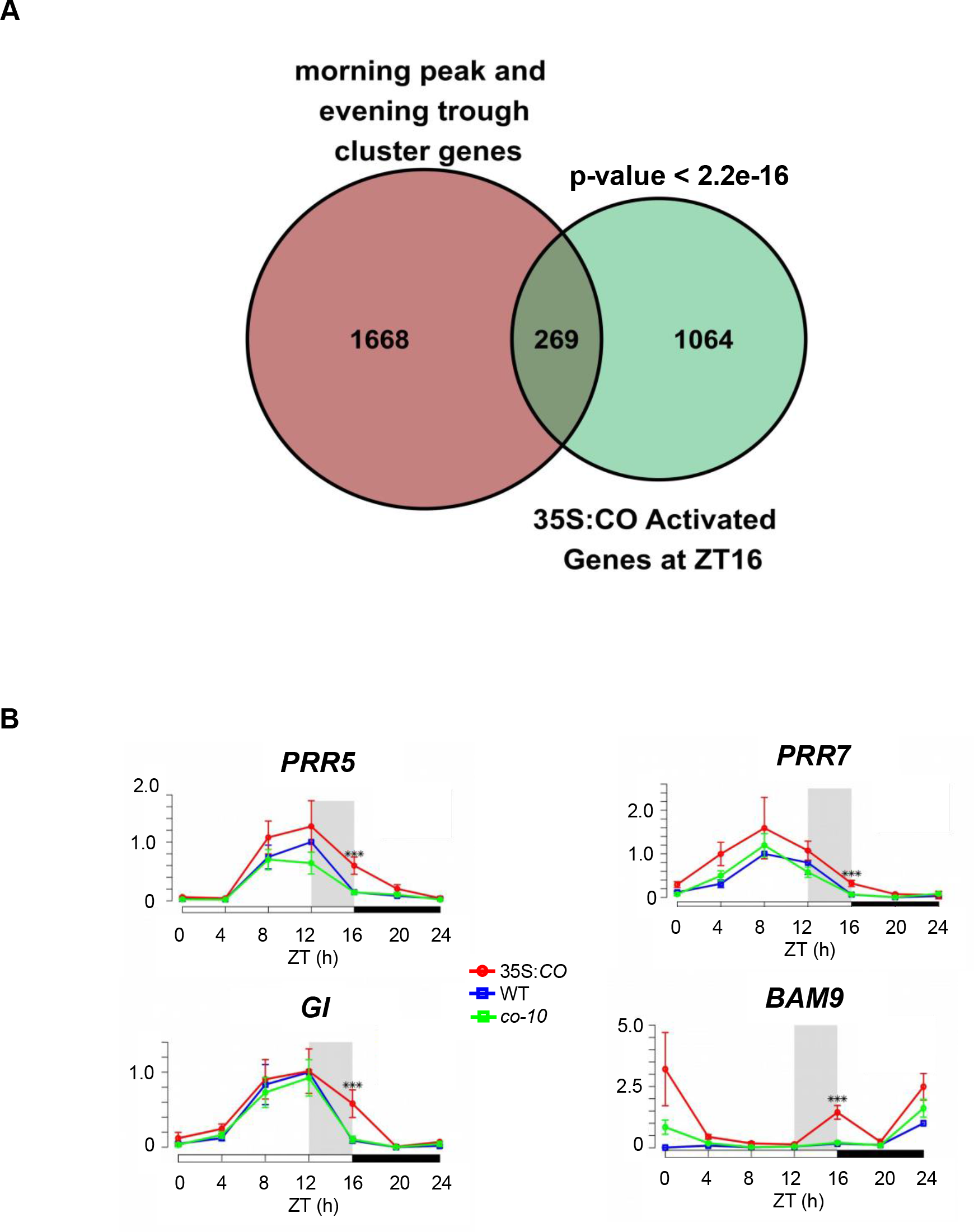
(A) Significant overlap between morning-phased cluster genes and the overexpressed genes in 35S:*CO* plants. The significance of the intersection was tested using Fisher’s Exact Test. (B) *PRR5*, *PRR7*, *GIGANTEA* and *BAM9* expression levels during a LD time course in Col-0, 35S:*CO* and *co-10* lines in whole seedlings, 7 DAG. The grey rectangle indicates the coincidence window in which the CO protein is stabilized. Error bars represent the standard deviation within two biological replicates. Three measurements were carried out for each biological replicate.

**Figure S4.**
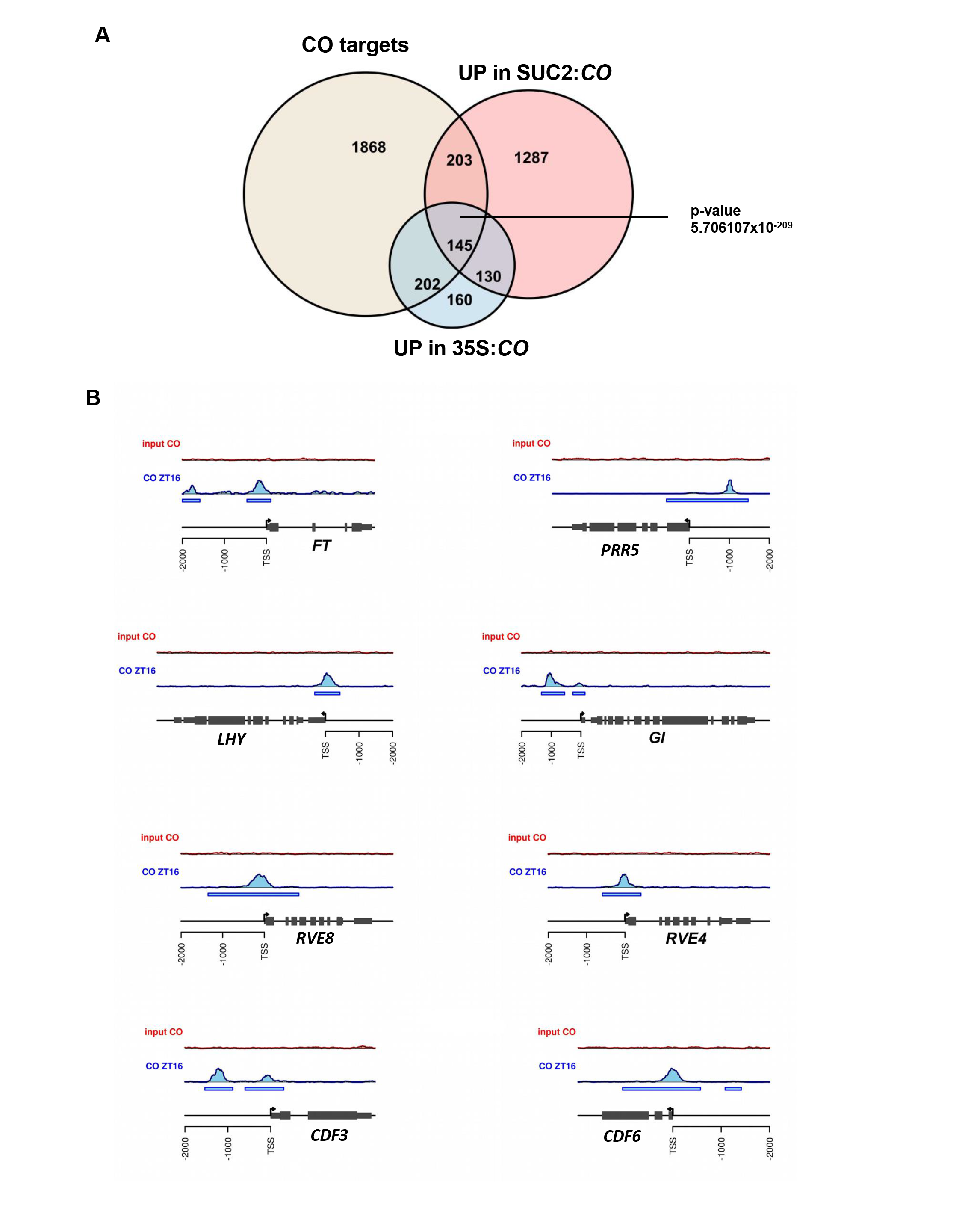
(A) Venn diagram of the CO targets from ChIP-seq data and upregulated genes in 35S:*CO* and SUC2:*CO* plants. (B) Genome browser views showing CO binding to the *FT*, *PRR5*, *LHY*, *GI*, *RVE8*, *RVE4*, *CDF3* and *CDF6* promoters.

**Figure S5.**
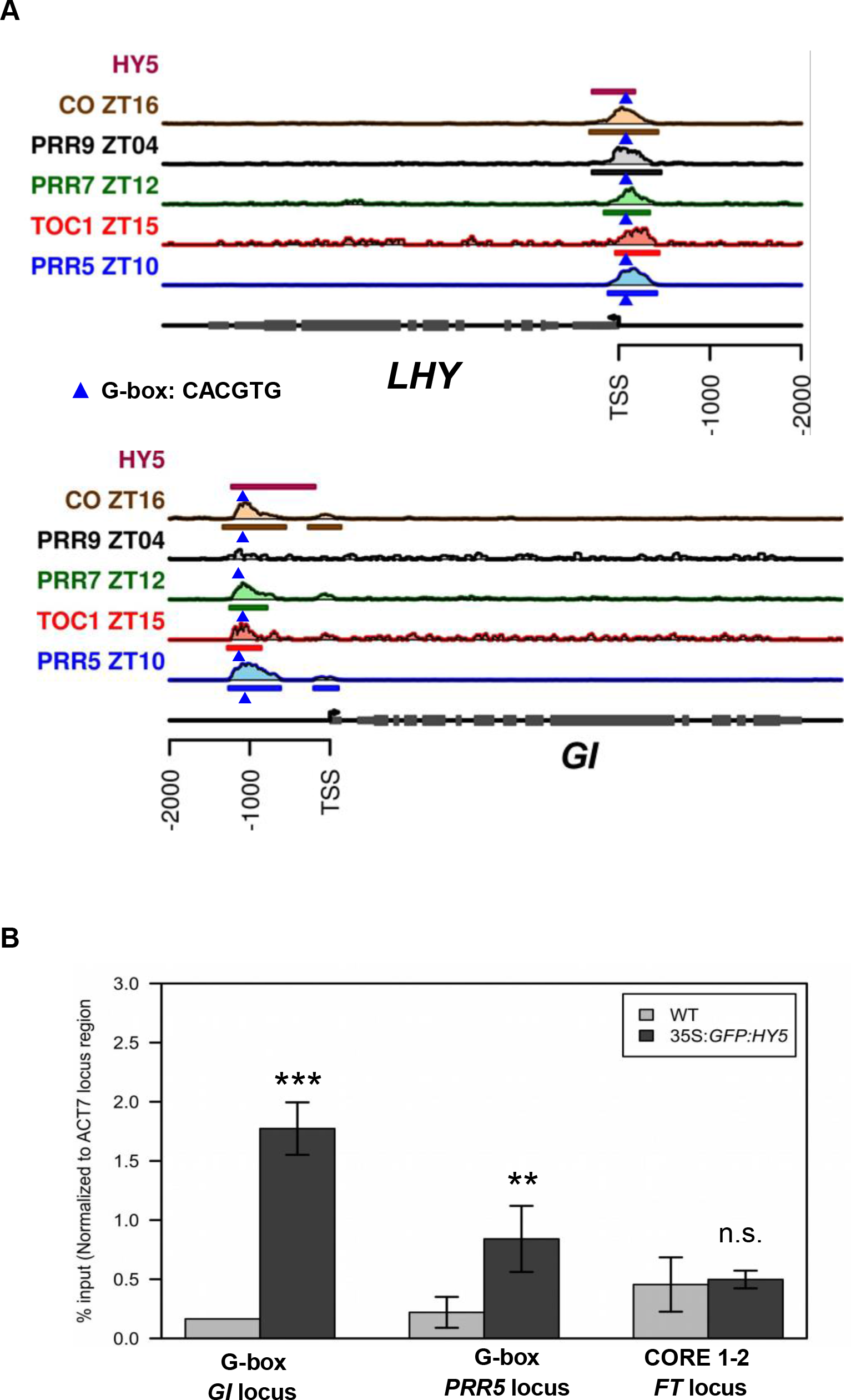
(A) Peak Visualizer views of CO, HY5, and PRRs (TOC1, PRR5, PRR7 and PRR9) binding to G-box-containing regions within the *LHY* and *GI* promoters. (B) HY5 binding to *PRR5*, *GI* and *FT* promoters. Here, 10-DAG-old plants grown under LD conditions were employed to perform ChIP assays. The bars represent the standard deviation between two biological replicates. Three technical replicates were measured for each biological replicate, asterisks indicate >0.99 significance.

**Figure S6.**
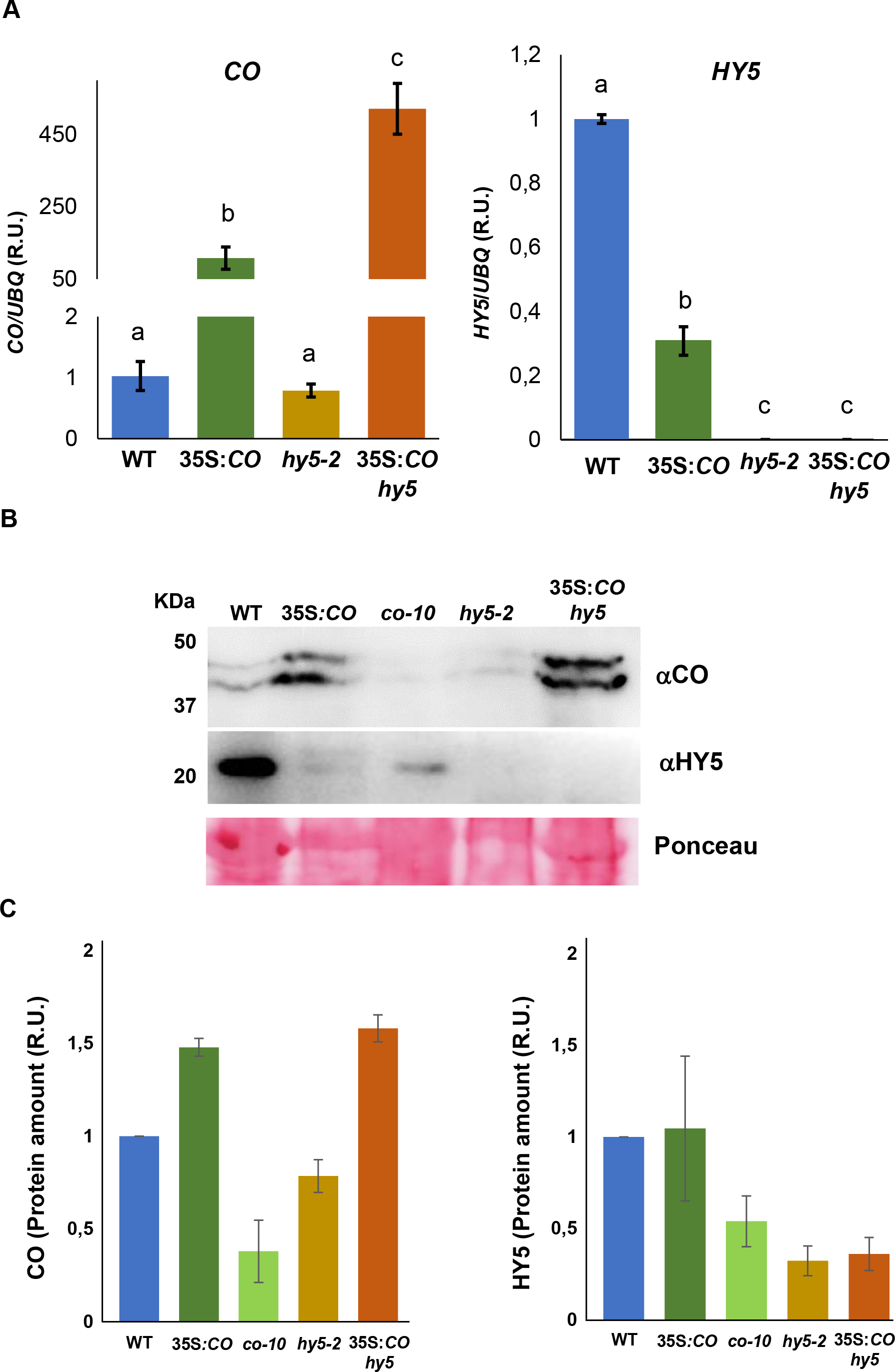
(A) *CO* and *HY5* expression levels measured by RT-qPCR in Col-0 (WT), 35S:*CO*, *hy5-2* and 35S:*CO hy5* lines. (B) Levels of CO and HY5 proteins in Col-0 (WT), 35S:*CO, co-10, hy5-2* and 35S:*CO hy5* plant extracts by Western blot using αCo and αHY5. Ponceau staining is shown as loading control. Molecular masses (kDa) are indicated on the left. Error bars indicate the standard deviation (s.d.) from three independent experiments. P < 0.05, one-way ANOVA and Tukey’s HSD.

**Figure S7.**
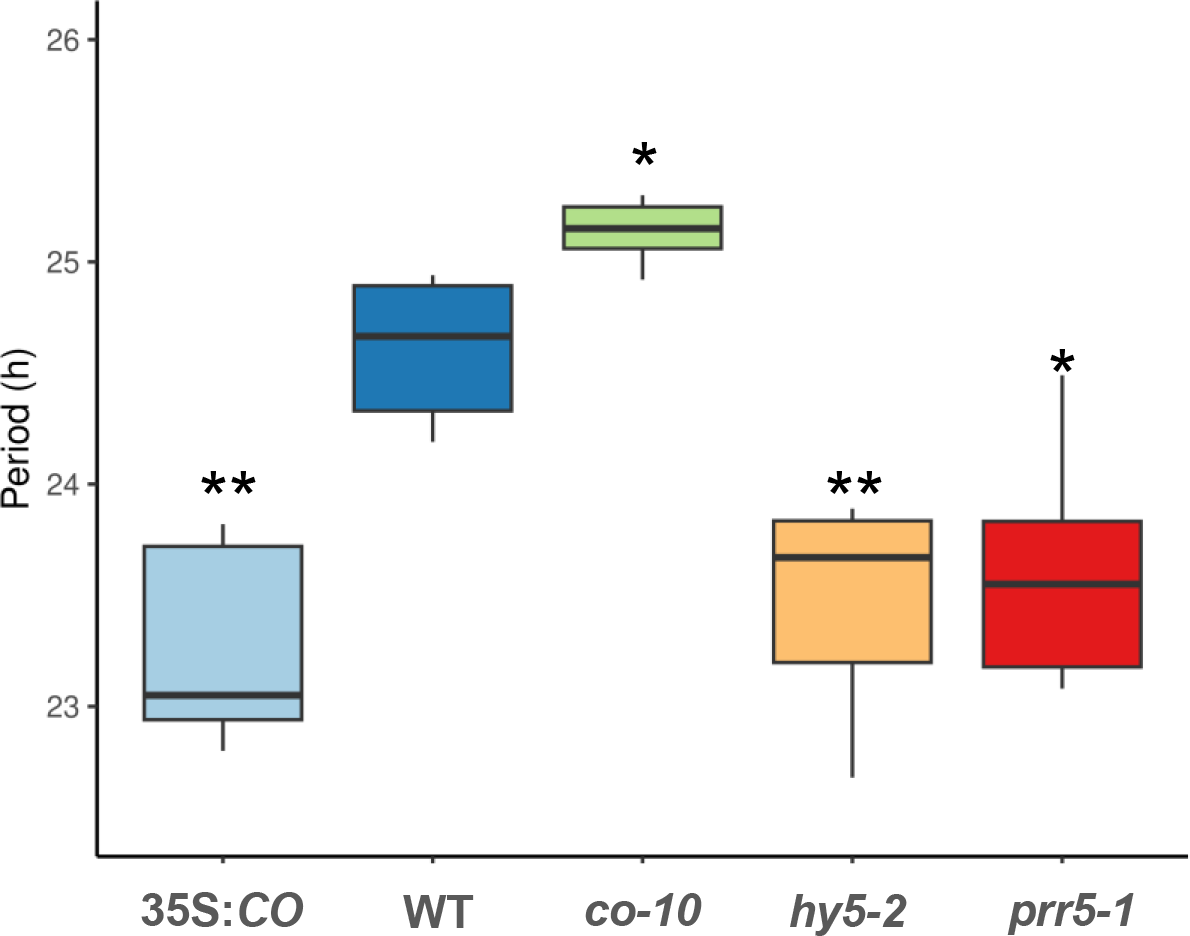
(A) Period of leaf movement in 35S:*CO*, Col-0 (WT), *co-10*, *hy5-2* and *prr5- 1* plants grown in LL, 7 DAG. * P < 0.05, ** P < 0.01.

## References

1. Abbas, N., Maurya, J. P., Senapati, D., Gangappa, S. N., and Chattopadhyaya, S. (2014). Arabidopsis CAM7 and HY5 physically interact and directly bind to the HY5 promoter to regulate its expression and thereby promote photomorphogenesis. Plant Cell 26:1036–1052.

2. Alon, U. (2007). Simplicity in biology. Nature 446:497.

3. An, H., Roussot, C., Suárez-López, P., Corbesier, L., Vincent, C., Piñeiro, M., Hepworth, S., Mouradov, A., Justin, S., Turnbull, C., et al. (2004). CONSTANS acts in the phloem to regulate a systemic signal that induces photoperiodic flowering of Arabidopsis. Development 131:3615–3626.

4. Ando, E., Ohnishi, M., Wang, Y., Matsushita, T., Watanabe, A., Hayashi, Y., Fujii, M., Ma, J. F., Inoue, S. I., and Kinoshita, T. (2013). TWIN SISTER OF FT, GIGANTEA, and CONSTANS have a positive but indirect effect on blue light-induced Stomatal opening in Arabidopsis. Plant Physiology 162:1529–1538.

5. Andrés, F., and Coupland, G. (2012). The genetic basis of flowering responses to seasonal cues. Nature Reviews Genetics 13:627–639.

6. Barabási, A.-L. (2015). Network Science.

7. Belda-Palazón, B., Ruiz, L., Martí, E., Tárraga, S., Tiburcio, A. F., Culiáñez, F., Farràs, R., Carrasco, P., and Ferrando, A. (2012). Aminopropyltransferases Involved in Polyamine Biosynthesis Localize Preferentially in the Nucleus of Plant Cells. PLoS ONE 7:e46907.

8. Bhagat, P. K., Verma, D., Sharma, D., and Sinha, A. K. (2021). HY5 and ABI5 transcription factors physically interact to fine tune light and ABA signaling in Arabidopsis. Plant Mol Biol 107:117–127.

9. Binkert, M., Kozma-Bognár, L., Terecskei, K., De Veylder, L., Nagy, F., and Ulm, R. (2014). UV-B-Responsive association of the Arabidopsis bZIP transcription factor ELONGATED HYPOCOTYL5 with target genes, including its own promoter. Plant Cell 26:4200–4213.

10. Blackman, B. K., and Michaels, S. D. (2010). Does CONSTANS act as a transcription factor or as a co-activator? The answer may be - yes. New Phytologist 187:1– 3.

11. Bowler, C., Benvenuto, G., Laflamme, P., Molino, D., Probst, A. V., Tariq, M., and Paszkowski, J. (2004). Chromatin techniques for plant cells. The Plant Journal 39:776–789.

12. Bursch, K., Toledo-Ortiz, G., Pireyre, M., Lohr, M., Braatz, C., and Johansson, H. (2020). Identification of BBX proteins as rate-limiting cofactors of HY5. Nat. Plants 6:921–928.

13. Chattopadhyay, S., Ang, L. H., Puente, P., Deng, X. W., and Wei, N. (1998). Arabidopsis bZIP protein HY5 directly interacts with light-responsive promoters in mediating light control of gene expression. Plant Cell 10:673–683.

14. Csardi, G. N. T. (2006). The igraph software package for complex network research. Inter J Complex Syst 1695.

15. Datta, S., Hettiarachchi, C., Johansson, H., and Holm, M. (2007). SALT TOLERANCE HOMOLOG2, a B-Box Protein in Arabidopsis That Activates Transcription and Positively Regulates Light-Mediated Development. Plant Cell 19:3242–3255.

16. de los Reyes, P., Romero-Campero, F. J., Teresa Ruiz, M., Romero, J. M., and Valverde, F. (2017). Evolution of daily gene co-expression patterns from algae to plants. Frontiers in Plant Science 8:1217.

17. De Mendoza, A., Sebé-Pedrós, A., Šestak, M. S., Matejčić, M., Torruella, G., Domazet-Lošo, T., and Ruiz-Trillo, I. (2013). Transcription factor evolution in eukaryotes and the assembly of the regulatory toolkit in multicellular lineages. Proceedings of the National Academy of Sciences of the United States of America 110:E4858–E4866.

18. Dobin, A., Davis, C. A., Schlesinger, F., Drenkow, J., Zaleski, C., Jha, S., Batut, P., Chaisson, M., and Gingeras, T. R. (2013). STAR: ultrafast universal RNA-seq aligner. Bioinformatics 29:15.

19. Endo, M., Tanigawa, Y., Murakami, T., Araki, T., and Nagatani, A. (2013). Phytochrome-dependent late-flowering accelerates flowering through physical interactions with phytochrome B and CONSTANS. Proceedings of the National Academy of Sciences of the United States of America 110:18017–18022.

20. Endo, M., Shimizu, H., Nohales, M. A., Araki, T., and Kay, S. A. (2014). Tissue-specific clocks in Arabidopsis show asymmetric coupling. Nature 515:419–422.

21. Fernández, V., Takahashi, Y., Le Gourrierec, J., and Coupland, G. (2016). Photoperiodic and thermosensory pathways interact through CONSTANS to promote flowering at high temperature under short days. The Plant Journal 86:426–440.

22. Gangappa, S. N., and Botto, J. F. (2016). The Multifaceted Roles of HY5 in Plant Growth and Development. Molecular Plant 9:1353–1365.

23. Gnesutta, N., Kumimoto, R. W., Swain, S., Chiara, M., Siriwardana, C., Horner, D. S., Holt, B. F., and Mantovani, R. (2017). CONSTANS imparts DNA sequence specificity to the histone fold NF-YB/NF-YC Dimer. Plant Cell 29:1516–1532.

24. Goldstein, M. L., Morris, S. A., and Yen, G. G. (2004). Problems with fitting to the power-law distribution. Eur. Phys. J. B 41:255–258.

25. Graeff, M., Straub, D., Eguen, T., Dolde, U., Rodrigues, V., Brandt, R., and Wenkel, S. (2016). MicroProtein-Mediated Recruitment of CONSTANS into a TOPLESS Trimeric Complex Represses Flowering in Arabidopsis. PLoS Genetics 12:e1005959.

26. Greenham, K., Lou, P., Remsen, S. E., Farid, H., and McClung, C. R. (2015). TRiP: Tracking Rhythms in Plants, an automated leaf movement analysis program for circadian period estimation. Plant Methods 11.

27. Gyuris, J., Golemis, E., Chertkov, H., and Brent, R. (1993). Cdi1, a human G1 and S phase protein phosphatase that associates with Cdk2. Cell 75:791–803.

28. Hajdu, A., Ádám, É., Sheerin, D. J., Dobos, O., Bernula, P., Hiltbrunner, A., Kozma-Bognár, L., and Nagy, F. (2015). High-level expression and phosphorylation of phytochrome B modulates flowering time in Arabidopsis. The Plant Journal 83:794–805.

29. Hajdu, A., Dobos, O., Domijan, M., Bálint, B., Nagy, I., Nagy, F., and Kozma-Bognár, L. (2018). ELONGATED HYPOCOTYL 5 mediates blue light signalling to the Arabidopsis circadian clock. Plant J 96:1242–1254.

30. Hayama, R., Sarid-Krebs, L., Richter, R., Fernández, V., Jang, S., and Coupland, G. (2017). PSEUDO RESPONSE REGULATORs stabilize CONSTANS protein to promote flowering in response to day length. The EMBO Journal 36:904–918.

31. Heinz, S., Benner, C., Spann, N., Bertolino, E., Lin, Y. C., Laslo, P., Cheng, J. X., Murre, C., Singh, H., and Glass, C. K. (2010). Simple combinations of lineage-determining transcription factors prime cis-regulatory elements required for macrophage and B cell identities. Molecular cell 38:576–89.

32. Huang, W., Pérez-García, P., Pokhilko, A., Millar, A. J., Antoshechkin, I., Riechmann, J. L., and Mas, P. (2012). Mapping the core of the Arabidopsis circadian clock defines the network structure of the oscillator. Science 335:75– 79.

33. Imaizumi, T., Schultz, T. F., Harmon, F. G., Ho, L. A., and Kay, S. A. (2005). Plant science: FKF1 F-box protein mediates cyclic degradation of a repressor of CONSTANS in Arabidopsis. Science 309:293–297.

34. Ito, S., Song, Y. H., Josephson-Day, A. R., Miller, R. J., Breton, G., Olmstead, R. G., and Imaizumi, T. (2012). FLOWERING BHLH transcriptional activators control expression of the photoperiodic flowering regulator CONSTANS in Arabidopsis. Proceedings of the National Academy of Sciences of the United States of America 109:3582–3587.

35. Jang, S., Marchal, V., Panigrahi, K. C. S., Wenkel, S., Soppe, W., Deng, X. W., Valverde, F., and Coupland, G. (2008). Arabidopsis COP1 shapes the temporal pattern of CO accumulation conferring a photoperiodic flowering response. EMBO Journal 27:1277–1288.

36. Jing, Y., Zhang, D., Wang, X., Tang, W., Wang, W., Huai, J., Xu, G., Chen, D., Li, Y., and Lin, R. (2013). Arabidopsis chromatin remodeling factor PICKLE interacts with transcription factor HY5 to regulate hypocotyl cell elongation. Plant Cell 25:242–256.

37. Jing, Y., Guo, Q., Zha, P., and Lin, R. (2019). The chromatin-remodelling factor PICKLE interacts with CONSTANS to promote flowering in Arabidopsis. Plant, Cell & Environment 42:2495–2507.

38. Job, N., Yadukrishnan, P., Bursch, K., Datta, S., and Johansson, H. (2018). Two B-Box Proteins Regulate Photomorphogenesis by Oppositely Modulating HY5 through their Diverse C-Terminal Domains. Plant Physiology 176:2963–2976.

39. Kinoshita, A., and Richter, R. (2020). Genetic and molecular basis of floral induction in Arabidopsis thaliana. J Exp Bot 71:2490–2504.

40. Langmead, B., Trapnell, C., Pop, M., and Salzberg, S. L. (2009). Ultrafast and memory-efficient alignment of short DNA sequences to the human genome. Genome biology 10:R25.

41. Lau, O. S., and Bergmann, D. C. (2015). MOBE-ChIP: a large-scale chromatin immunoprecipitation assay for cell type-specific studies. The Plant journal : for cell and molecular biology 84:443–50.

42. Laubinger, S., Marchal, V., Gentillhomme, J., Wenkel, S., Adrian, J., Jang, S., Kulajta, C., Braun, H., Coupland, G., and Hoecker, U. (2006). Arabidopsis SPA proteins regulate photoperiodic flowering and interact with the floral inducer CONSTANS to regulate its stability. Development 133:3213–3222.

43. Lazaro, A., Valverde, F., Piñeiro, M., and Jarillo, J. A. (2012). The Arabidopsis E3 ubiquitin ligase HOS1 negatively regulates CONSTANS abundance in the photoperiodic control of flowering. Plant Cell 24:982–999.

44. Lazaro, A., Mouriz, A., Piñeiro, M., and Jarillo, J. A. (2015). Red Light-Mediated Degradation of CONSTANS by the E3 Ubiquitin Ligase HOS1 Regulates Photoperiodic Flowering in Arabidopsis. The Plant cell 27:2437–54.

45. Lee, J., He, K., Stolc, V., Lee, H., Figueroa, P., Gao, Y., Tongprasit, W., Zhao, H., Lee, I., and Deng, X. W. (2007). Analysis of transcription factor HY5 genomic binding sites revealed its hierarchical role in light regulation of development. The Plant cell 19:731–49.

46. Liao, Y., Smyth, G. K., and Shi, W. (2014). featureCounts: an efficient general purpose program for assigning sequence reads to genomic features. *Bioinformatics (Oxford*, England) 30:923–930.

47. Liu, L. J., Zhang, Y. C., Li, Q. H., Sang, Y., Mao, J., Lian, H. L., Wang, L., and Yang, H. Q. (2008). COP1-mediated ubiquitination of CONSTANS is implicated in cryptochrome regulation of flowering in Arabidopsis. Plant Cell 20:292–306.

48. Liu, B., Zuo, Z., Liu, H., Liu, X., and Lin, C. (2011). Arabidopsis cryptochrome 1 interacts with SPA1 to suppress COP1 activity in response to blue light. Genes and Development 25:1029–1034.

49. Liu, T., Carlsson, J., Takeuchi, T., Newton, L., and Farré, E. M. (2013). Direct regulation of abiotic responses by the Arabidopsis circadian clock component PRR7. The Plant journal : for cell and molecular biology 76:101–14.

50. Liu, T. L., Newton, L., Liu, M.-J., Shiu, S.-H., and Farré, E. M. (2016). A G-Box-Like Motif Is Necessary for Transcriptional Regulation by Circadian Pseudo-Response Regulators in Arabidopsis. Plant Physiology 170:528–539.

51. Martín, G., Rovira, A., Veciana, N., Soy, J., Toledo-Ortiz, G., Gommers, C. M. M., Boix, M., Henriques, R., Minguet, E. G., Alabadí, D., et al. (2018). Circadian Waves of Transcriptional Repression Shape PIF-Regulated Photoperiod-Responsive Growth in Arabidopsis. Current Biology 28:311–318.e5.

52. McClung, C. R. (2006). Plant circadian rhythms. The Plant cell 18:792–803.

53. McClung, C. R. (2014). Wheels within wheels: New transcriptional feedback loops in the Arabidopsis circadian clock. F1000Prime Reports 6.

54. Mellis, I. A., and Raj, A. (2015). Half dozen of one, six billion of the other: What can small- and large-scale molecular systems biology learn from one another? Genome Research 25:1466–1472.

55. Michael, T. P., Salomé, P. A., Yu, H. J., Spencer, T. R., Sharp, E. L., McPeek, M. A., Alonso, J. M., Ecker, J. R., and McClung, C. R. (2003). Enhanced Fitness Conferred by Naturally Occurring Variation in the Circadian Clock. Science 302:1049–1053.

56. Milo, R., Shen-Orr, S., Itzkovitz, S., Kashtan, N., Chklovskii, D., and Alon, U. (2002). Network motifs: Simple building blocks of complex networks. Science 298:824–827.

57. Mizoguchi, T., Wright, L., Fujiwara, S., Cremer, F., Lee, K., Onouchi, H., Mouradov, A., Fowler, S., Kamada, H., Putterill, J., et al. (2005). Distinct roles of GIGANTEA in promoting flowering and regulating circadian rhythms in Arabidopsis. Plant Cell 17:2255–2270.

58. Nakamichi, N., Kita, M., Niinuma, K., Ito, S., Yamashino, T., Mizoguchi, T., and Mizuno, T. (2007). Arabidopsis Clock-Associated Pseudo-Response Regulators PRR9, PRR7 and PRR5 Coordinately and Positively Regulate Flowering Time Through the Canonical CONSTANS-Dependent Photoperiodic Pathway. Plant and Cell Physiology 48:822–832.

59. Nakamichi, N., Kiba, T., Henriques, R., Mizuno, T., Chua, N.-H., and Sakakibara, H. (2010). PSEUDO-RESPONSE REGULATORS 9, 7, and 5 are transcriptional repressors in the Arabidopsis circadian clock. The Plant cell 22:594–605.

60. Nakamichi, N., Kiba, T., Kamioka, M., Suzuki, T., Yamashino, T., Higashiyama, T., Sakakibara, H., and Mizuno, T. (2012). Transcriptional repressor PRR5 directly regulates clock-output pathways. Proceedings of the National Academy of Sciences of the United States of America 109:17123–8.

61. Nguyen, K. T., Park, J., Park, E., Lee, I., and Choi, G. (2015). The Arabidopsis RING Domain Protein BOI Inhibits Flowering via CO-dependent and CO-independent Mechanisms. Molecular Plant 8:1725–1736.

62. Onouchi, H., Igeño, M. I., Périlleux, C., Graves, K., and Coupland, G. (2000). Mutagenesis of Plants Overexpressing CONSTANS Demonstrates Novel Interactions among Arabidopsis Flowering-Time Genes. The Plant Cell 12:885.

63. Ortiz-Marchena, M. I., Albi, T., Lucas-Reina, E., Said, F. E., Romero-Campero, F. J., Cano, B., Teresa Ruiz, M., Romero, J. M., and Valverde, F. (2014). Photoperiodic control of carbon distribution during the floral transition in Arabidopsis. Plant Cell 26:565–584.

64. Osterlund, M. T., Hardtke, C. S., Ning, W., and Deng, X. W. (2000). Targeted destabilization of HY5 during light-regulated development of Arabidopsis. Nature 405:462–466.

65. Parsons, R., Parsons, R., Garner, N., Oster, H., and Rawashdeh, O. (2020). CircaCompare: A method to estimate and statistically support differences in mesor, amplitude and phase, between circadian rhythms. Bioinformatics 36:1208–1212.

66. Piñeiro, M., and Jarillo, J. A. (2013). Ubiquitination in the control of photoperiodic flowering. Plant Science 198:98–109.

67. Putterill, J., Robson, F., Lee, K., Simon, R., and Coupland, G. (1995). The CONSTANS gene of arabidopsis promotes flowering and encodes a protein showing similarities to zinc finger transcription factors. Cell 80:847–857.

68. Ritchie, M. E., Phipson, B., Wu, D., Hu, Y., Law, C. W., Shi, W., and Smyth, G. K. (2015). limma powers differential expression analyses for RNA-sequencing and microarray studies. Nucleic Acids Research 43:e47–e47.

69. Robinson, M. D., McCarthy, D. J., and Smyth, G. K. (2010). edgeR: a Bioconductor package for differential expression analysis of digital gene expression data. Bioinformatics 26:139–140.

70. Romero, J. M., and Valverde, F. (2009). Evolutionarily conserved photoperiod mechanisms in plants: When did plant photoperiodic signaling appear? Plant Signaling and Behavior 4:642–644.

71. Romero-Campero, F. J., Lucas-Reina, E., Said, F. E., Romero, J. M., and Valverde, F. (2013). A contribution to the study of plant development evolution based on gene co-expression networks. Frontiers in plant science 4:291.

72. Rugnone, M. L., Faigón Soverna, A., Sanchez, S. E., Schlaen, R. G., Hernando, C. E., Seymour, D. K., Mancini, E., Chernomoretz, A., Weigel, D., Más, P., et al. (2013). LNK genes integrate light and clock signaling networks at the core of the Arabidopsis oscillator. Proceedings of the National Academy of Sciences of the United States of America 110:12120–5.

73. Salmon-Divon, M., Dvinge, H., Tammoja, K., and Bertone, P. (2010). PeakAnalyzer: Genome-wide annotation of chromatin binding and modification loci. BMC Bioinformatics 2010 *11*:1 11:1–12.

74. Sanchez, S. E., and Kay, S. A. (2016). The Plant Circadian Clock: From a Simple Timekeeper to a Complex Developmental Manager. Cold Spring Harbor perspectives in biology 8:a027748.

75. Sawa, M., Nusinow, D. A., Kay, S. A., and Imaizumi, T. (2007). FKF1 and GIGANTEA complex formation is required for day-length measurement in Arabidopsis. Science 318:261–265.

76. Schindelin, J., Arganda-Carreras, I., Frise, E., Kaynig, V., Longair, M., Pietzsch, T., Preibisch, S., Rueden, C., Saalfeld, S., Schmid, B., et al. (2012). Fiji: An open-source platform for biological-image analysis. Nature Methods 9:676–682.

77. Serrano-Bueno, G., Said, F. E., de Los Reyes, P., Lucas-Reina, E. I., Ortiz-Marchena, M. I., Romero, J. M., and Valverde, F. (2020). CONSTANS- FKBP12 interaction contributes to modulation of photoperiodic flowering in Arabidopsis. Plant J 101:1287–1302.

78. Serrano-Bueno, G., Sánchez de Medina Hernández, V., and Valverde, F. (2021). Photoperiodic Signaling and Senescence, an Ancient Solution to a Modern Problem? Front Plant Sci 12:634393.

79. Serrano-Bueno, G., de los Reyes, P., Chini, A., Ferreras-Garrucho, G., Sánchez de Medina-Hernández, V., Boter, M., Solano, R., and Valverde, F. (2022). Regulation of floral senescence in Arabidopsis by coordinated action of CONSTANS and jasmonate signaling. Molecular Plant Advance Access published September 2022, doi:10.1016/J.MOLP.2022.09.017.

80. Sheerin, D. J., Menon, C., Oven-Krockhaus, S. Z., Enderle, B., Zhu, L., Johnen, P., Schleifenbaum, F., Stierhof, Y. D., Huq, E., and Hiltbrunner, A. (2015). Light-activated phytochrome A and B interact with members of the SPA family to promote photomorphogenesis in arabidopsis by reorganizing the COP1/SPA complex. Plant Cell 27:189–201.

81. Shen, C., Liu, H., Guan, Z., Yan, J., Zheng, T., Yan, W., Wu, C., Zhang, Q., Yin, P., and Xing, Y. (2020). Structural Insight into DNA Recognition by CCT/NF-YB/YC Complexes in Plant Photoperiodic Flowering. The Plant cell 32:3469–3484.

82. Shen-Orr, S. S., Milo, R., Mangan, S., and Alon, U. (2002). Network motifs in the transcriptional regulation network of Escherichia coli. Nature genetics 31:64– 68.

83. Shim, J. S., Kubota, A., and Imaizumi, T. (2017). Circadian clock and photoperiodic flowering in arabidopsis: CONSTANS is a Hub for Signal integration. Plant Physiology 173:5–15.

84. Shyu, Y. J., Suarez, C. D., and Hu, C. D. (2008). Visualization of ternary complexes in living cells by using a BiFC-based FRET assay. Nature Protocols 3:1693– 1702.

85. Smoot, M. E., Ono, K., Ruscheinski, J., Wang, P.-L., and Ideker, T. (2011). Cytoscape 2.8: new features for data integration and network visualization. *Bioinformatics (Oxford*, England) 27:431–2.

86. Song, Y. H., Smith, R. W., To, B. J., Millar, A. J., and Imaizumi, T. (2012). FKF1 conveys timing information for CONSTANS stabilization in photoperiodic flowering. Science 336:1045–1049.

87. Song, Y. H., Estrada, D. A., Johnson, R. S., Kim, S. K., Lee, S. Y., MacCoss, M. J., and Imaizumi, T. (2014). Distinct roles of FKF1, GIGANTEA, and ZEITLUPE proteins in the regulation of constans stability in Arabidopsis photoperiodic flowering. Proceedings of the National Academy of Sciences of the United States of America 111:17672–17677.

88. Stone, L., Simberloff, D., and Artzy-Randrup, Y. (2019). Network motifs and their origins. PLOS Computational Biology 15:e1006749.

89. Strayer, C., Oyama, T., Schultz, T. F., Raman, R., Somers, D. E., Mas, P., Panda, S., Kreps, J. A., and Kay, S. A. (2000). Cloning of the Arabidopsis clock gene TOC1, an autoregulatory response regulator homolog. Science 289:768–771.

90. Suárez-López, P., Wheatley, K., Robson, F., Onouchi, H., Valverde, F., and Coupland, G. (2001). CONSTANS mediates between the circadian clock and the control of flowering in Arabidopsis. Nature 410:1116–1120.

91. Thaben, P. F., and Westermark, P. O. (2014). Detecting rhythms in time series with RAIN. Journal of biological rhythms 29:391–400.

92. Tiwari, S. B., Shen, Y., Chang, H.-C., Hou, Y., Harris, A., Ma, S. F., McPartland, M., Hymus, G. J., Adam, L., Marion, C., et al. (2010). The flowering time regulator CONSTANS is recruited to the FLOWERING LOCUS T promoter via a unique cis-element. New Phytologist 187:57–66.

93. Tong, A. H. Y., Lesage, G., Bader, G. D., Ding, H., Xu, H., Xin, X., Young, J., Berriz, G. F., Brost, R. L., Chang, M., et al. (2004). Global Mapping of the Yeast Genetic Interaction Network. Science 303:808–813.

94. Valverde, F. (2011). CONSTANS and the evolutionary origin of photoperiodic timing of flowering. Journal of experimental botany 62:2453–63.

95. Valverde, F., Mouradov, A., Soppe, W., Ravenscroft, D., Samach, A., and Coupland, G. (2004). Photoreceptor Regulation of CONSTANS Protein in Photoperiodic Flowering. Science 303:1003–1006.

96. Wagner, A., and Fell, D. A. (2001). The small world inside large metabolic networks. Proceedings of the Royal Society of London. Series B: Biological Sciences 268:1803–1810.

97. Wang, Z.Y., and Tobin, E.M. (1998). Constitutive expression of the *CIRCADIAN CLOCK ASSOCIATED 1* (*CCA1*) gene disrupts circadian rhythms and suppresses its own expression. Cell 93: 1207–1217.

98. Wang, C. Q., Guthrie, C., Sarmast, M. K., and Dehesh, K. (2014). BBX19 interacts with CONSTANS to repress FLOWERING LOCUS T transcription, defining a flowering time checkpoint in Arabidopsis. Plant Cell 26:3589–3602.

99. Wang, H., Pan, J., Li, Y., Lou, D., Hu, Y., and Yu, D. (2016). The DELLA-CONSTANS transcription factor cascade integrates gibberellic acid and photoperiod signaling to regulate flowering. Plant Physiology 172:479–488.

100. Wenkel, S., Turck, F., Singer, K., Gissot, L., Le Gourrierec, J., Samach, A., and Coupland, G. (2006). CONSTANS and the CCAAT box binding complex share a functionally important domain and interact to regulate flowering of Arabidopsis. The Plant cell 18:2971–84.

101. Wheeler, D. L., Barrett, T., Benson, D. A., Bryant, S. H., Canese, K., Church, D. M., DiCuccio, M., Edgar, R., Federhen, S., Helmberg, W., et al. (2005). Database resources of the National Center for Biotechnology Information. Nucleic acids research 33:D39–45.

102. Wu, T., Hu, E., Xu, S., Chen, M., Guo, P., Dai, Z., Feng, T., Zhou, L., Tang, W., Zhan, L., et al. (2021). clusterProfiler 4.0: A universal enrichment tool for interpreting omics data. The Innovation 2:100141.

103. Xu, F., Li, T., Xu, P.-B., Li, L., Du, S.-S., Lian, H.-L., and Yang, H.-Q. (2016). DELLA proteins physically interact with CONSTANS to regulate flowering under long days in *Arabidopsis*. FEBS Letters 590:541–549.

104. Yook, S.-H., Oltvai, Z. N., and Barabási, A.-L. (2004). Functional and topological characterization of protein interaction networks. PROTEOMICS 4:928–942.

105. Young, M.W., and Kay, S.A. (2001). Time zones: A comparative genetics of circadian clocks. Nat. Rev. Genet. 2: 702–715.

106. Young, H. S., Cheol, M. Y., An, P. H., Seong, H. K., Hee, J. J., Su, Y. S., Hye, J. K., Yun, D. J., Chae, O. L., Jeong, D. B., et al. (2008). DNA-binding study identifies C-box and hybrid C/G-box or C/A-box motifs as high-affinity binding sites for STF1 and long hypocotyl5 proteins. Plant Physiology 146:1862–1877.

107. Yu, G., Wang, L.-G., and He, Q.-Y. (2015). ChIPseeker: an R/Bioconductor package for ChIP peak annotation, comparison and visualization. Bioinformatics 31:2382–2383.

108. Yu, B., He, X., Tang, Y., Chen, Z., Zhou, L., Li, X., Zhang, C., Huang, X., Yang, Y., Zhang, W., et al. (2023). Photoperiod controls plant seed size in a CONSTANS- dependent manner. Nat. Plants 9:343–354.

109. Zhang, Y., Liu, T., Meyer, C. A., Eeckhoute, J., Johnson, D. S., Bernstein, B. E., Nusbaum, C., Myers, R. M., Brown, M., Li, W., et al. (2008). Model-based analysis of ChIP-Seq (MACS). Genome biology 9:R137.

110. Zhang, H., He, H., Wang, X., Wang, X., Yang, X., Li, L., and Deng, X. W. (2011). Genome-wide mapping of the HY5-mediated genenetworks in Arabidopsis that involve both transcriptional and post-transcriptional regulation. The Plant Journal 65:346–358.

111. Zhang, J. D., Ruschhaupt, M., and Biczok, R. (2016). ddCt method for qRT–PCR data analysis Advance Access published 2016.

112. Zhu, L. J., Gazin, C., Lawson, N. D., Pagès, H., Lin, S. M., Lapointe, D. S., and Green, M. R. (2010). ChIPpeakAnno: a Bioconductor package to annotate ChIP-seq and ChIP-chip data. BMC Bioinformatics 2010 11:1 11:1–10.

113. Zielinski, T., Moore, A. M., Troup, E., Halliday, K. J., and Millar, A. J. (2014). Strengths and Limitations of Period Estimation Methods for Circadian Data. PLOS ONE 9:e96462.

114. Zuo, Z., Liu, H., Liu, B., Liu, X., and Lin, C. (2011). Blue light-dependent interaction of CRY2 with SPA1 regulates COP1 activity and floral initiation in arabidopsis. Current Biology 21:841–847.

