## Supplemental Table 9 for "CONSTANS alters the circadian clock in *Arabidopsis thaliana*"

**Table S9**. Primers used in this study.

Primers used to measure amplicon enrichment in ChIP-QPCR experiments.

| **Region** | **Primer sequence** | **Amplicon length** |
| --- | --- | --- |
| TSS *ACT2* | GTAACATTGTGCTCAGTGGTGG | 286 |
|  | CTCGGCCTTGGAGATCCACATC |  |
| Random | CAACGAGATTTGGGGTTAG | 259 |
|  | TTGAATGCAGTCCGATTGTCC |  |
| CORE 1/2 *FT* promoter | GTGGCTACCAAGTGGGAGAT | 199 |
|  | TAACTCGGGTCGGTGAAATC |  |
| G-box *PRR5* promoter | AGGTGAAAGACTGTGTCAGATAA | 106 |
|  | TGTTGGGTGGGATTAGATTAGAG |  |
| G-box *GI* promoter | ATCACGAATCGTATGGAGATCATTA | 110 |
|  | CAATCAGAGTGGAGCAAGAGAT |  |

Primers used to measure gene expression by RT-qPCR

| **Gene** | **Primer sequence** | **Amplicon length** |
| --- | --- | --- |
| *UBQ10* | GAAGTTCAATGTTTCGTTTCATGT | 123 |
|  | GGATTATACAAGGCCCCAAAA |  |
| *PRR5* | GCAATCTCTTCAACGAGAAGCCGC | 138 |
|  | CGAACGAATTGGCCTTTGATTCG |  |
| *PRR7* | GGAAGTGGTAGCGGAAACTTGG | 110 |
|  | CGTACCTTCTTTCGGAAGCACC |  |
| *GI* | CCAGCAACAATACGGTGCC | 158 |
|  | CGATAGGACGGACTATTCATTCCG |  |
| *BAM9* | CTTGATGGGAAGACTCCTATGG | 98 |
|  | GTGATTCCCGTGATTGTGTTG |  |
| *CO* | CCAATGGACAGAGAAGCCAGG | 175 |
|  | GCATCGTGTTGAACCCTTGC |  |
| *HY5* | GCTGAAGAGGTTGTTGAGGA | 101 |
|  | TCTCCAAGTCTTTCACTCTGTTT |  |
