## Supplemental Table 10 for "CONSTANS alters the circadian clock in *Arabidopsis thaliana*"

**Table S10**. Transcription factors ChIP-seq data used to build the transcriptional network CircadianFloralNet. Accession numbers of the study, samples and number of target genes are indicated.

| **Transcription factor (Accession number)** | **Samples** | **Number of target genes** |
| --- | --- | --- |
| SOC1 - At2g45660  (GSE45846) | SRR822347 | 1328 |
|  | SRR822348 |  |
|  | SRR822349 |  |
|  | SRR822350 |  |
|  | SRR822351 |  |
|  | SRR822352 |  |
| PIF4 - At2g43010  (GSE43284) | SRR643944 | 1038 |
|  | SRR643945 |  |
|  | SRR643946 |  |
|  | SRR643947 |  |
|  | SRR643948 |  |
|  | SRR643949 |  |
| FHY1 - At2g37678  (GSE58083) | SRR1302274 | 6539 |
|  | SRR1302275 |  |
|  | SRR1302276 |  |
|  | SRR1302277 |  |
| APETALA1 - At1g69120  (GSE46987) | SRR1042993 | 2420 |
|  | SRR851697 |  |
|  | SRR851698 |  |
|  | SRR851699 |  |
|  | SRR851700 |  |
|  | SRR851701 |  |
|  | SRR851702 |  |
| SEPALLATA3 - At1g24260  (GSE46987) | SRR1042994 | 5709 |
|  | SRR1042995 |  |
|  | SRR851693 |  |
|  | SRR851694 |  |
|  | SRR851695 |  |
|  | SRR851696 |  |
|  | SRR851702 |  |
| PHYA - At1g09570  (GSE48769) | SRR931764 | 7911 |
|  | SRR931765 |  |
|  | SRR931766 |  |
|  | SRR931767 |  |
| FLM - At1g77080  (GSE48082) | SRR908234 | 3627 |
|  | SRR908235 |  |
|  | SRR908236 |  |
|  | SRR908237 |  |
|  | SRR908238 |  |
|  | SRR908239 |  |
|  | SRR908240 |  |
|  | SRR908241 |  |
|  | SRR908242 |  |
|  | SRR908243 |  |
|  | SRR908244 |  |
|  | SRR908245 |  |
|  | SRR908246 |  |
|  | SRR908247 |  |
| SVP - At2g22540  (GSE33120) | SRR354187 | 149 |
|  | SRR354188 |  |
|  | SRR354189 |  |
|  | SRR354190 |  |
|  | SRR354191 |  |
|  | SRR354192 |  |
| APETALA3 - At3g54340  (GSE38358) | SRR502859 | 8501 |
|  | SRR502860 |  |
| PISTILLATA - At5g20240  (GSE38358) | SRR502857 | 6316 |
|  | SRR502858 |  |
|  | SRR502860 |  |
| FLC - At5g10140  (*) | SRR116764 | 4 |
|  | SRR116765 |  |
| TOC1 - At5g61380  (GSE35952) | SRR411113 | 274 |
|  | SRR411114 |  |
| AGAMOUS - At4g18960  (GSE45938) | SRR824573 | 2295 |
|  | SRR824574 |  |
|  | SRR824575 |  |
|  | SRR824576 |  |
| LEAFY - At5g61850  (GSE24568) | SRR070382 | 4598 |
|  | SRR070383 |  |
|  | SRR070384 |  |
|  | SRR070385 |  |
| PIF4 - At2g43010  (GSE43284) | SRR643944 | 1038 |
|  | SRR643945 |  |
|  | SRR643946 |  |
|  | SRR643947 |  |
|  | SRR643948 |  |
|  | SRR643949 |  |
| ARF6 - At1g30330  (GSE51770) | SRR1019434 | 1984 |
|  | SRR1019435 |  |
| JAG - At1g68480  (GSE51537) | SRR1015049 | 292 |
|  | SRR1015050 |  |
|  | SRR1015051 |  |
|  | SRR1015052 |  |
|  | SRR1015053 |  |
|  | SRR1015054 |  |
| PIF5 - At3g59060  (GSE35059) | SRR398909 | 1789 |
|  | SRR398910 |  |
| PIF3 - At1g09530  (GSE39215) | SRR520249 | 229 |
|  | SRR520250 |  |
|  | SRR520251 |  |
|  | SRR520252 |  |
|  | SRR520253 |  |
|  | SRR520254 |  |
|  | SRR520255 |  |
|  | SRR520256 |  |
|  | SRR520257 |  |
|  | SRR520258 |  |
|  | SRR520259 |  |
|  | SRR520260 |  |
| PIF1 - At2g20180  (GSE43283) | SRR643938 | 3190 |
|  | SRR643939 |  |
|  | SRR643940 |  |
|  | SRR643941 |  |
|  | SRR643942 |  |
|  | SRR643943 |  |
| SPEECHLESS - At5g53210  (GSE57497) | SRR1282200 | 6027 |
|  | SRR1282201 |  |
| HBI1 - At2g18300  (GSE53099) | SRR1044949 | 15 |
|  | SRR1044950 |  |
| AL5 - At5g20510  (GSE56706) | SRR1232232 | 24 |
|  | SRR1232233 |  |
| KAN1 - At5g16560  (GSE48081) | SRR908249 | 6881 |
|  | SRR908250 |  |
|  | SRR908251 |  |
|  | SRR908252 |  |
| APETALA 2 - At4g36920  (GSE21301) | SRR040045 | 1572 |
|  | SRR040046 |  |
|  | SRR040047 |  |
|  | SRR040048 |  |
|  | SRR040049 |  |
|  | SRR040050 |  |
|  | SRR040051 |  |
| ERF115 - At5g07310  (GSE48793) | SRR931836 | 1266 |
|  | SRR931837 |  |
| IBH1 - At4g30410  (GSE51120) | SRR1001911 | 59 |
|  | SRR1001912 |  |
| LHY - At1g01060  (GSE52175) | SRR1027034 | 86 |
|  | SRR1027035 |  |
| SPL7 – At5g18830 (GSE45213) | SRR776581 | 3 |
|  | SRR776582 |  |
| CCA1 - At2g46830  (GSE70533) | SRR2087595 | 5485 |
|  | SRR2087596 |  |
|  | SRR2087597 |  |
|  | SRR2087598 |  |
|  | SRR2087599 |  |
|  | SRR2087560 |  |
|  | SRR2087561 |  |
| PRR9 - At2g46790  (GSE71397) | SRR2131056 | 137 |
|  | SRR2131057 |  |
| PRR7 - At5g02810  (GSE49282) | SRR943788 | 1873 |
|  | SRR943789 |  |
| FHY3 - At3g22170  (GSE30713) | SRR309188 | 1802 |
|  | SRR309189 |  |
