## Supplemental Table 11 for "CONSTANS alters the circadian clock in *Arabidopsis thaliana*"

**Table S11**. RNA-seq datasets analyzed in this study. The accession numbers, samples and conditions are indicated.

| **Experiment** | **Condition** | **Sample** |
| --- | --- | --- |
| Genome-wide analysis of LINK1 and LINK2 effects on the Arabidopsis transcriptome (GSE43865) | Long days WT, ZT2. | SRR653561 |
|  |  | SRR653562 |
|  |  | SRR653563 |
|  | Long days WT, ZT6. | SRR653564 |
|  |  | SRR653565 |
|  |  | SRR653566 |
|  | Long days WT, ZT10 | SRR653567 |
|  |  | SRR653568 |
|  |  | SRR653569 |
|  | Long days WT, ZT14 | SRR653570 |
|  |  | SRR653571 |
|  |  | SRR653572 |
|  | Long days WT, ZT18 | SRR653573 |
|  |  | SRR653574 |
|  |  | SRR653575 |
|  | Long days WT, ZT22 | SRR653576 |
|  |  | SRR653577 |
|  |  | SRR653578 |
| Genome-wide analysis of *co* mutation effects on the Arabidopsis transcriptome (GSE205675) | Continuous light, *co-10* | SRR19579114 |
|  |  | SRR19579113 |
|  | Continuous light, Col-0 | SRR19579110 |
|  |  | SRR19579109 |
| Genome-wide analysis of CO effects on the Arabidopsis transcriptome (GSE236178) | Col-0 (WT), ZT14. | GSM7519351 |
|  |  | GSM7519352 |
|  |  | GSM7519353 |
|  | SUC2:*CO*, ZT14. | GSM7519354 |
|  |  | GSM7519355 |
|  |  | GSM7519356 |
|  | 35S:*CO,* ZT14. | GSM7519348 |
|  |  | GSM7519349 |
|  |  | GSM7519350 |
